# TLR4 Modulates Simvastatin’s Impact on HDL Cholesterol and Glycemic Control

**DOI:** 10.1101/2025.02.08.637204

**Authors:** Xiao Tian, Peixiang Zhang

## Abstract

Statins lower cardiovascular risk by inhibiting HMG-CoA reductase, with their efficacy and adverse effects linked to SREBP-regulated cholesterol metabolism. Nutritional states— particularly the alternating fasting and feeding cycles, rather than intrinsic circadian rhythms— exert tight control over hepatic lipid metabolism via SREBP-mediated regulation of cholesterol and fatty acid synthesis. We exploited the natural diurnal oscillations in food intake in mice to determine how nutritional state modulates simvastatin’s metabolic actions. When administered during fasting, simvastatin impaired glucose homeostasis through SREBP-2–driven autophagy that enhanced PPARα activity, whereas simvastatin given during feeding reduced HDL cholesterol and improved glucose metabolism. Mechanistically, a feeding-induced surge in lipopolysaccharide (LPS) suppressed hepatic oxysterol synthesis, thereby enabling simvastatin to further deplete oxysterols and inhibit activation of LXR and SREBP-1c—key regulators of PPARα signaling under both postprandial and fasting conditions. This cascade resulted in decreased HDL cholesterol levels and improved glycemic control. Moreover, TLR4 deficiency disrupted the LPS–simvastatin interplay on LXR/SREBP-1c/PPARα signaling, reversing the HDL-lowering and glucose-improving effects seen during feeding. Together, these results identify feeding-induced LPS–TLR4 signaling as a critical determinant of simvastatin’s metabolic effects and suggest strategies to optimize statin therapy based on nutritional state.

## INTRODUCTION

Atherosclerotic cardiovascular disease (ASCVD) remains the leading cause of mortality worldwide. By inhibiting the rate-limiting enzyme HMG-CoA reductase in cholesterol biosynthesis, statins effectively lower low-density lipoprotein cholesterol (LDL-C) levels and reduce ASCVD risk (Strandberg et al., 2024). Randomized controlled trials consistently show that a 1 mmol/L reduction in LDL-C decreases the incidence of myocardial infarction and ischemic stroke by approximately 25% (Baigent et al., 2010; Fulcher et al., 2015). Statins also elevate high-density lipoprotein cholesterol (HDL-C), promoting reverse cholesterol transport, mitigating oxidative stress, and enhancing endothelial function. Nonetheless, recent meta-analyses have identified a dose-dependent relationship between statin use and an increased risk (10–36%) of new-onset diabetes, highlighting the need to balance efficacy with potential adverse effects (Cholesterol Treatment Trialists’ Collaboration, 2024).

Growing evidence suggests that the timing of statin administration and food intake significantly affects both the effectiveness and safety of statins, particularly those with shorter plasma half-lives (e.g., simvastatin, fluvastatin, lovastatin) (Awad et al., 2017; Izquierdo-Palomares et al., 2016; Kim et al., 2013; Martin et al., 2002; Whitfield et al., 2000; McLean et al., 2018; Alakhali et al., 2018; Maqsood et al., 2022; Wang et al., 2023). While one systematic review and meta-analysis showed that evening administration of short half-life statins yielded greater LDL-C reductions (Awad et al., 2017), a Cochrane review (Izquierdo-Palomares et al., 2016) reported no significant lipid profile differences across different dosing times. These discrepancies imply that unexamined factors—especially feeding and fasting states—may play pivotal roles in mediating statin efficacy and tolerability (Stokkan et al., 2001; Vollmers et al., 2009; Greco et al., 2021).

Recent human and animal studies strongly indicate that feeding and fasting cycles, rather than intrinsic circadian rhythms, primarily drive hepatic lipid and cholesterol metabolism via sterol regulatory element-binding proteins (SREBPs) (Stokkan et al., 2001; Vollmers et al., 2009; Greco et al., 2021). SREBPs are central regulators of hepatic lipid metabolism, governing the expression of cholesterol and fatty acid synthesis genes. Intake of meals rich in carbohydrates and low in fat robustly activates nuclear SREBPs and their downstream targets, including HMG-CoA reductase, the rate-limiting enzyme in cholesterol synthesis (Horton et al., 1998). By contrast, fasting markedly reduces nuclear SREBP-1 and SREBP-2 levels and suppresses their target genes, with refeeding restoring their activity (Horton et al., 1998). Notably, these feeding-driven effects persist in models lacking functional circadian clocks, underscoring the supremacy of feeding cues over circadian regulation (Stokkan et al., 2001; Vollmers et al., 2009; Greco et al., 2021).

The feeding–fasting cycle precisely modulates SREBP activity and the expression of fatty acid and cholesterol synthesis genes (e.g., fatty acid synthetase, HMG-CoA reductase), aligning with shifts in hepatic cholesterol production rates (Popják et al., 1981; Kopito et al., 1982; Horton et al., 1998; Stokkan et al., 2001; Vollmers et al., 2009; Greco et al., 2021). Feeding can enhance hepatic HMG-CoA reductase activity by over fourfold, mirroring increased cholesterol synthesis (Kopito et al., 1982). Although plasma mevalonate—a downstream product of HMG-CoA reductase—has been proposed as a surrogate for cholesterol synthesis, it peaks about 12 hours after feeding in both rodents and humans, largely owing to renal uptake, urinary excretion, and nighttime reductions in urinary output (Nakamoto et al., 2021). While this delayed plasma mevalonate peak may limit its utility as a direct marker of real-time cholesterol synthesis (Kopito et al., 1982; McNamara et al., 1985; Nakamoto et al., 2021), these oscillations in hepatic HMG-CoA reductase activity and plasma mevalonate are abolished without feeding, emphasizing the dominant influence of feeding over circadian rhythms (Popják et al., 1981; Parker et al., 1984).

Given this body of evidence, we hypothesized that feeding behavior modulates statin efficacy and side effects by regulating hepatic SREBP-driven lipid and cholesterol metabolism. To test this hypothesis, we designed two simvastatin administration strategies in mice: (1) mixing simvastatin into their food during the active feeding phase, and (2) orally gavaging simvastatin during the fasting (sleep) phase. Our results show that feeding-phase simvastatin lowers HDL and improves glucose homeostasis, whereas fasting-phase simvastatin elevates HDL and impairs glucose tolerance. Further, we identify a feeding-driven surge of lipopolysaccharide (LPS) signaling via Toll-like receptor 4 (TLR4) as a key mechanism impacting the liver X receptor (LXR)/SREBP-1c/peroxisome proliferator-activated receptor alpha (PPARα) pathways, thereby influencing simvastatin’s effects on HDL and glucose metabolism.

## RESULTS

### Fasting Versus Feeding: Divergent Effects of Simvastatin on HDL and Glucose Metabolism

Under standard chow conditions, mice consume approximately 80% of their food during the dark (active) feeding phase (ZT12–ZT24) and minimally during the light (rest) fasting phase (ZT0– ZT12) (Turek et al., 2005; Brown et al., 2021). Capitalizing on this natural diurnal rhythm, we investigated whether fasting versus feeding states would differentially modulate simvastatin’s lipid and glucose regulatory effects in A/J mice over five weeks. We used two dosing strategies: simvastatin incorporated into chow (0.1 g/kg food weight; formulation D11060903i, Research Diets), ensuring drug ingestion during the active feeding phase, and oral gavage of an equivalent dose (16 mg/kg body weight) at 10:00 AM (ZT04), coinciding with the light-phase fasting period (Figure 1A, B). A typical 25-gram mouse consumes approximately 3–5 grams of food daily, corresponding to about 16 mg/kg body weight, which aligns with the orally gavaged dose. This approach preserved normal feeding rhythms and allowed a direct comparison of simvastatin’s effects when co-administered with food versus during fasting. Mice were housed on a 12-hour light/dark cycle (lights on at 6:00 AM [ZT0] and off at 6:00 PM [ZT12]) with ad libitum access to food and water. The 16 mg/kg dose was chosen to approximate an 80 mg human equivalent daily dose (Zhang et al., 2024), maintaining translational relevance.

**Figure 1.**
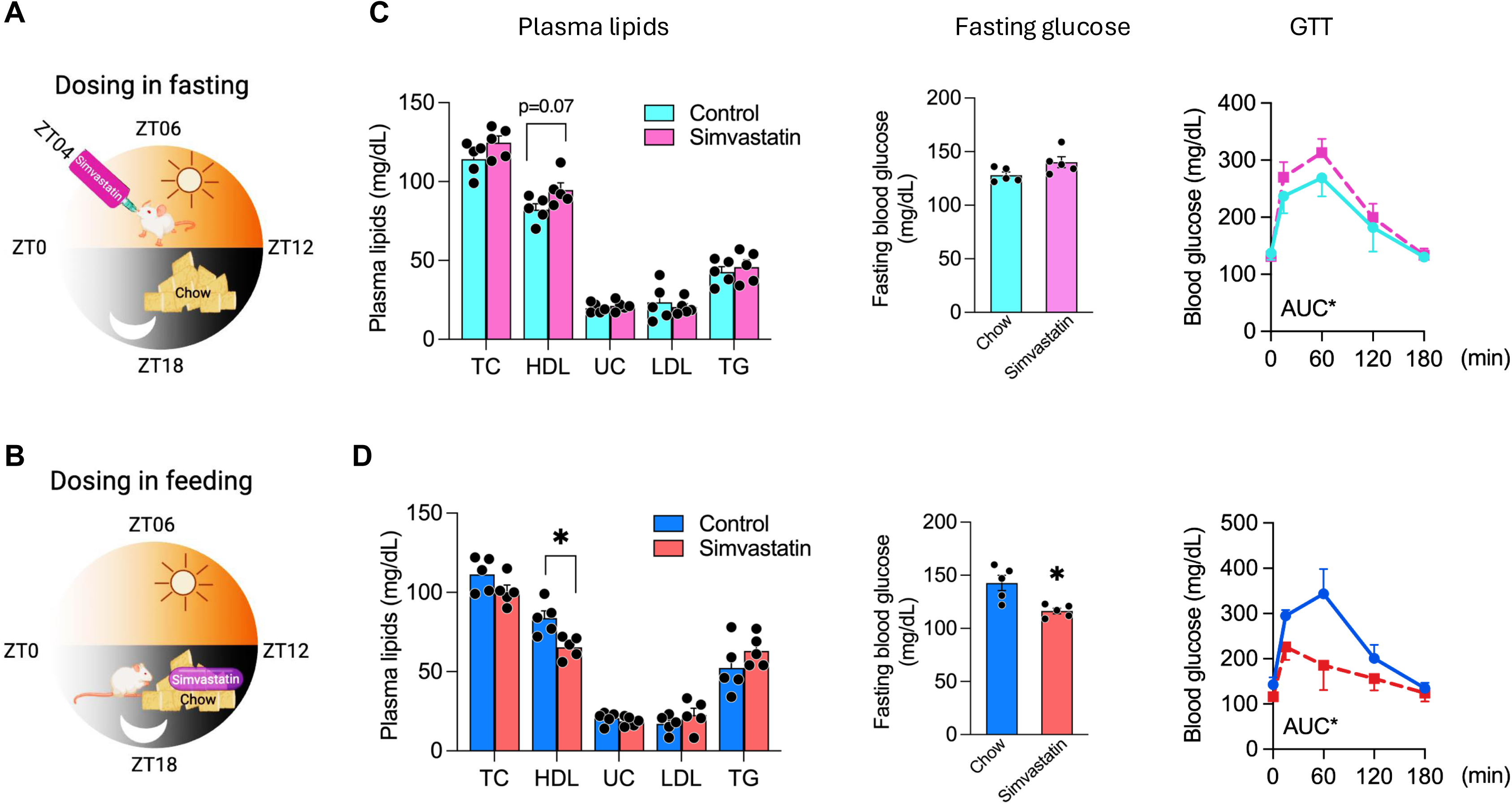
Differential Effects of Simvastatin Administration during Fasting or Feeding on HDL Cholesterol and Glucose Metabolism. **(A, B)** Schematic representations of the 5-week simvastatin treatment regimens. Mice were housed on a 12-hr light/dark cycle (lights on at 06:00, ZT0; lights off at 18:00, ZT12). In **(A)**, simvastatin was delivered by oral gavage at ZT04 (10:00 AM) during the fasting phase; in **(B)**, simvastatin was incorporated into standard chow, allowing consumption during the feeding phase. **(C)** Plasma lipid and glucose parameters following fasting-phase simvastatin administration, including high-density lipoprotein cholesterol (HDL-C), total cholesterol (TC), unesterified cholesterol (UC), triglycerides (TG), fasting glucose levels, and area under the curve (AUC) for glucose tolerance tests (GTTs). **(D)** Plasma lipid and glucose parameters following feeding-phase simvastatin administration, measured as in **(C)**. Data are presented as mean ± SEM (n = 5 per group), Statistical significance was determined as *p < 0.05.

After five weeks, we collected plasma and tissues at two time points to capture the drug’s effects in distinct metabolic states: the fasting-state group (ZT09, 3:00 PM), five hours after the last oral gavage, and the feeding-state group (ZT05, 11:00 AM), five hours after the active eating period. Mice receiving simvastatin by gavage during fasting showed a modest increase in HDL-C (p = 0.07), with no significant changes in total cholesterol (TC), unesterified cholesterol (UC), or triglycerides (TG) (Figure 1C). Low-density lipoprotein cholesterol (LDL-C) also remained unchanged, consistent with the characteristically low LDL-C in rodents (which lack cholesteryl ester transfer protein, CETP) (Steenbergen et al., 2010). Notably, fasting-phase simvastatin markedly impaired glucose metabolism, as evidenced by higher glucose area under the curve (AUC) values in glucose tolerance tests (Figure 1C). In contrast, co-administration of simvastatin with food significantly lowered HDL-C levels and improved glucose homeostasis, indicated by reduced fasting blood glucose and enhanced glucose tolerance (Figure 1D). Together, these data indicate that administering simvastatin during fasting versus feeding exerts opposing effects on HDL and glucose metabolism.

### Fasting and Feeding States Govern Simvastatin’s Regulation of PPARα

To determine how fasting and feeding states modulate simvastatin’s effects on HDL-C, we measured the expression of genes involved in HDL synthesis (apolipoprotein A1 [*Apoa1*], *Apoa2*, *Apom*, and ATP-binding cassette subfamily A member 1 [*Abca1*]), HDL maturation (lecithin-cholesterol acyltransferase [*Lcat*] and phospholipid transfer protein [*Pltp*]), and HDL clearance (scavenger receptor class B type 1 [*Scarb1*], encoding SR-B1) (Paththinige et al., 2017). When simvastatin was administered during fasting, *Pltp* expression increased significantly, and *Apoa1* showed a modest increase (p < 0.1), whereas *Apoa2*, *Apom*, *Abca1*, *Lcat*, and *Scarb1* remained unchanged (Figure 2A). By contrast, simvastatin administration during feeding suppressed both *Apoa1* and *Apoa2*. These findings illustrate how fasting and feeding states critically influence simvastatin’s impact on HDL metabolism, particularly through effects on apolipoproteins and phospholipid transfer protein (Figures 1C–D and 2A).

**Figure 2.**
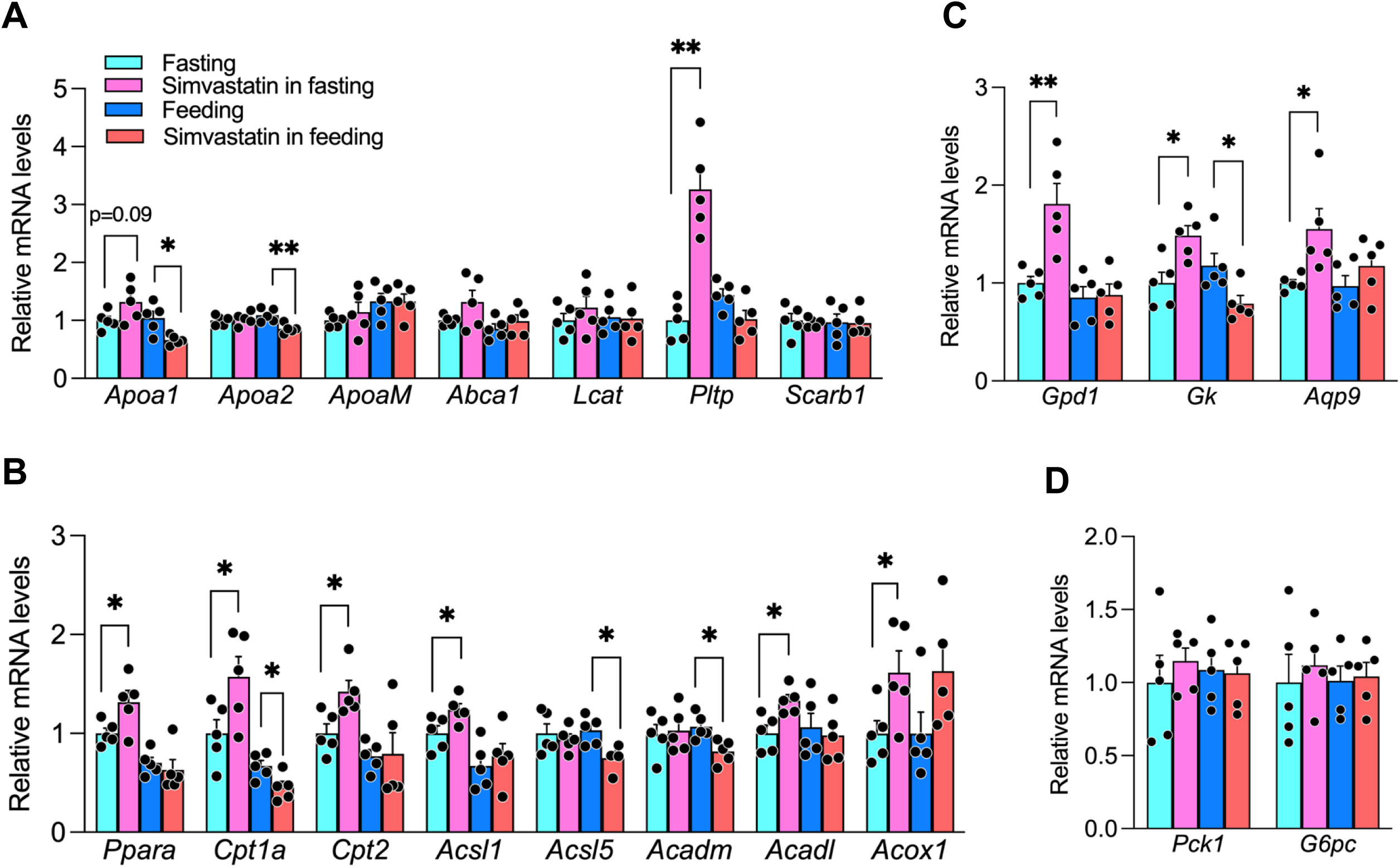
Hepatic Gene Expression Modulation by Simvastatin Administration in Fasting and Feeding Phases. **(A)** Relative mRNA levels of hepatic high-density lipoprotein (HDL)-related genes. Liver samples were collected 5 hr after the final simvastatin administration—ZT09 (3:00 PM) for the fasting group and ZT05 (11:00 AM) for the feeding group. **(B)** Hepatic expression of peroxisome proliferator-activated receptor alpha (PPARα) and its downstream targets. **(C)** Gene expression profiling of gluconeogenesis-related genes. **(D)** Expression levels of pyruvate/lactate pathway genes. Each bar represents the mean ± SEM (n = 5 per group). All qRT-PCR data were normalized to *Hprt*. Statistical significance is indicated by *p < 0.05; **p < 0.01.

Because *Apoa1*, *Apoa2*, and *Pltp* are regulated by PPARα ligands such as fibrates and given that statins can activate PPARα to induce Apoa1 expression in human cells, we hypothesized that simvastatin in fasting or feeding states would differentially modulate PPARα’s regulation of apolipoproteins and *Pltp*, thereby affecting HDL-C levels (Bouly et al., 2001; Martin et al., 2001; Han et al., 2006). Indeed, simvastatin administered during fasting significantly upregulated PPARα and its direct targets, including carnitine palmitoyltransferase 1A (*Cpt1a*), carnitine palmitoyltransferase 2 (*Cpt2*), acyl-CoA synthetase long chain family member 1 (*Acsl1*), acyl-CoA dehydrogenase long chain (*Acadl*), and acyl-CoA oxidase 1 (*Acox1*). Conversely, simvastatin co-administered with food reduced the expression of key PPARα targets, such as *Cpt1a,* acyl-CoA dehydrogenase medium chain (*Acadm*), and acyl-CoA synthetase long chain family member 5 (*Acsl5*) (Figure 2B). In conjunction with the *Apoa1*, *Apoa2*, and *Pltp* findings (Figure 2A), these results underscore the critical influence of fasting and feeding states on simvastatin-mediated PPARα activity.

In addition to regulating lipid metabolism, PPARα plays a major role in glucose homeostasis, particularly by driving glycerol-fueled gluconeogenesis (Patsouris et al., 2004). To assess how fasting or feeding states affect gluconeogenesis-related pathways, we examined genes involved in these processes. Fasting-phase simvastatin treatment significantly enhanced the expression of glycerol-3-phosphate dehydrogenase 1 (*Gpd1*), glycerol kinase (*Gk*), and aquaporin 9 (*Aqp9*), a glycerol transporter, whereas feeding-phase administration decreased their expression (Figure 2C). In contrast, simvastatin—whether administered during fasting or feeding—did not alter the expression of genes in the pyruvate/lactate pathway (phosphoenolpyruvate carboxykinase 1 [*Pck1*] and glucose-6-phosphatase catalytic subunit [*G6pc*]) (Figure 2D). Thus, simvastatin administration in fasting or feeding states differentially affects PPARα activation, influencing glycerol-driven gluconeogenesis and contributing to the distinct glycemic responses observed between fasting and fed conditions (Figure 1C–D).

### SREBP-2-Driven Autophagy Links Simvastatin to PPARα Activation

We confirmed that simvastatin mimics the fasting state in vitro. Serum-starving Huh-7 cells overnight and then exposing them to increasing doses of simvastatin led to a dose-dependent induction of P*LTP*, *APOA1*, and *GPD1* (Figure 3A), consistent with the fasting-phase effects observed in vivo (Figure 2A). Although free fatty acids (FFAs) derived from fasting-induced adipose tissue lipolysis can activate PPARα (Bideyan et al., 2021), treating Huh-7 cells with oleic acid alone, however, did not increase *PLTP*, *APOA1*, or *GPD1* expression, suggesting that that FFAs require additional signals to activate PPARα. Interestingly, when oleic acid was combined with simvastatin, *APOA1* expression was markedly elevated compared to simvastatin alone (Figure 3B), indicating that simvastatin can potentiate FFA mediated PPARα activation.

**Figure 3.**
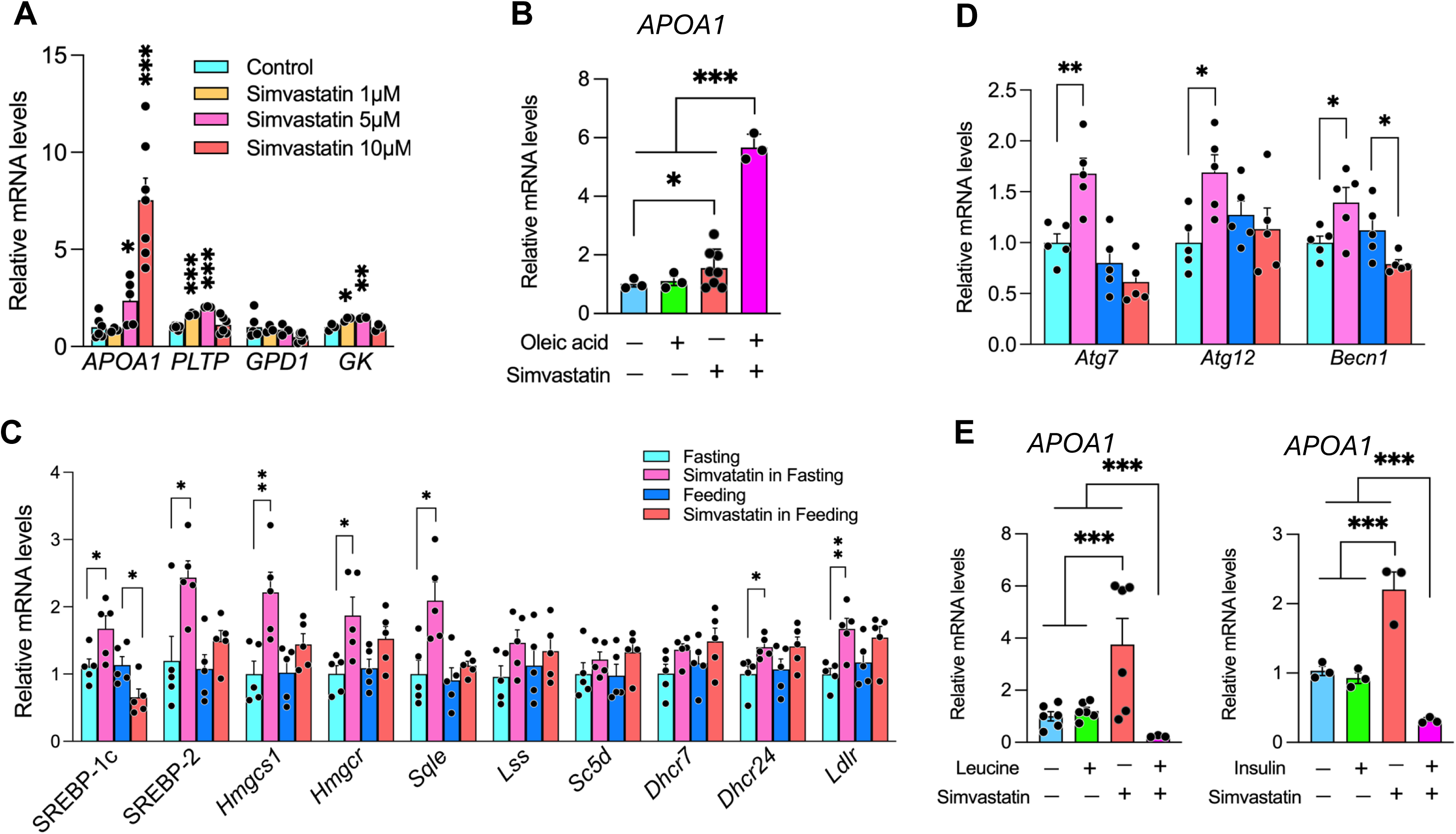
Simvastatin Administration during Fasting Activates SREBP-2 and Autophagy and Augments FFA-Induced APOA1 Expression. **(A)** Huh7 cells cultured under serum-depleted conditions were treated with increasing concentrations of simvastatin (1, 5, or 10 μM) for 24 hr. *APOA1*, *PLTP*, and *GK* mRNA levels were determined by qPCR and normalized to the geometric mean of *HPRT* and *B2M*. Data are shown as fold change relative to untreated controls. **(B)** Simvastatin (10 μM) enhances sodium oleate (50 μM)–stimulated APOA1 expression in serum-depleted Huh7 cells. mRNA levels were measured by qPCR and normalized to the geometric mean of *HPRT* and *B2M*. **(C)** Fasting-phase simvastatin treatment upregulates SREBP-2 target genes in mouse liver. Animals were sacrificed 5 hr after the final simvastatin dose (ZT09, 3:00 PM), and hepatic mRNA levels were quantified by qPCR and normalized to *Hprt*. **(D)** Expression of the autophagy-related genes in liver tissue following fasting-phase simvastatin treatment, measured by qPCR and normalized to *Hprt*. **(E)** Autophagy inhibition (with leucine and insulin) attenuates simvastatin-mediated enhancement of oleate-induced PPARα and *APOA1* expression in Huh7 cells. Gene expression was quantified by qPCR and normalized to the geometric mean of *HPRT* and *B2M*. Data are presented as mean ± SEM. Individual data points are overlaid on bar graphs (n values indicated). Statistical significance: *p < 0.05; **p < 0.01; ***p < 0.001. All qPCR analyses were performed in triplicate.

Autophagy enhances PPARα activation by degrading corepressors such as NCoR1 and HDAC3 (Saito et al., 2019; Iershov et al., 2019), and SREBP-2 transactivation can trigger autophagy (Seo et al., 2011). In liver tissues from fasting mice, simvastatin markedly increased SREBP-2 expression and its downstream cholesterol synthesis targets by 1- to 3-fold (Figure 3C), concomitant with upregulation of key autophagy-related genes including *Atg7, Atg12*, and *Becn1* (Figure 3D). These findings indicate that simvastatin activates SREBP-2–dependent autophagy during fasting, which likely amplifies FFA-mediated PPARα activation. To directly test whether autophagy is essential for this synergistic effect, serum-starved Huh-7 cells were treated with simvastatin in the presence of leucine or insulin—both of which activate mTORC1 and inhibit autophagy (Fernandes et al., 2024). Under these autophagy-inhibitory conditions, simvastatin-induced *APOA1* expression was abolished, and the potentiating simvastatin induction of PPARα targets (*PLTP*, *APOA1*, and *GPD1*) was significantly attenuated (Figure 3E). Collectively, these data support a model in which simvastatin triggers SREBP-2–dependent autophagy during fasting, thereby amplifying FFA-mediated PPARα activation and ultimately modulating HDL levels and hepatic glucose metabolism in vivo.

### Postprandial LPS Undermines Simvastatin’s Activation of PPARα

Co-administration of simvastatin with food suppressed PPARα activation, contrasting with the enhanced activation observed when simvastatin was given during fasting (Figures 2B–C). To identify the food intake–associated factors that could suppress PPARα, we assessed the effects of various nutrients and hormones on simvastatin-induced APOA1 under serum-replete conditions (10% FBS). Neither leucine nor insulin influenced simvastatin-induced APOA1 (Figure 4A), and neither oleic acid nor high glucose levels altered simvastatin’s effects under the same culture conditions (data not shown). These observations suggest that, under serum-replete conditions, common dietary nutrients and insulin exert minimal influence on simvastatin-mediated PPARα activation.

**Figure 4.**
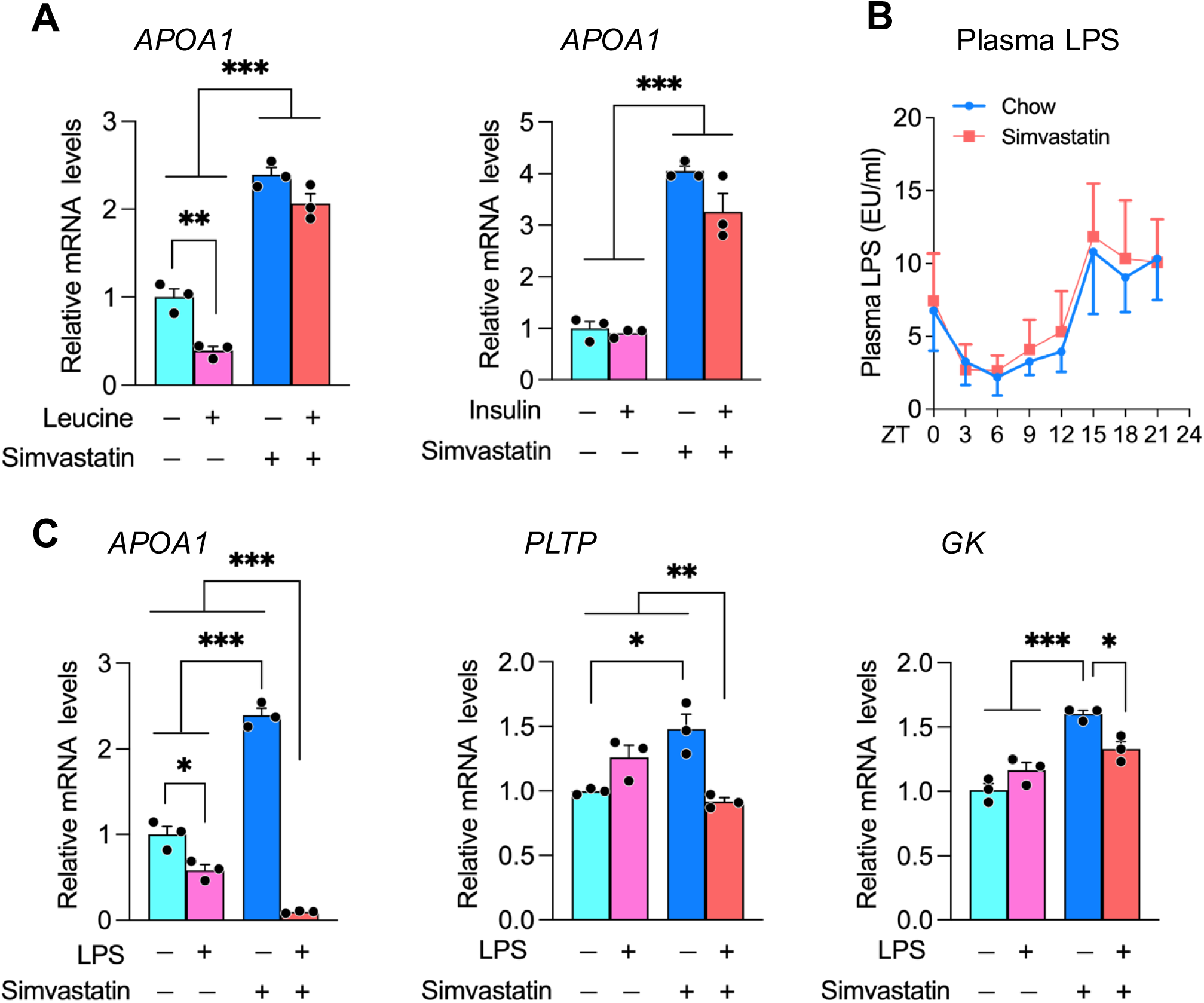
Interplay Between Lipopolysaccharide (LPS) and Simvastatin in Regulating APOA1, PLTP, and GK Expression. **(A)** Impact of leucine and insulin co-treatment with simvastatin on *APOA1* expression in Huh7 cells. Cells were incubated under the indicated conditions, and *APOA1* mRNA levels were measured by qPCR. **(B)** Plasma LPS levels measured in mice during the light (fasting) phase (ZT0–12) and dark (feeding) phase (ZT12–24), illustrating alignment with feeding patterns. **(C)** Effect of LPS co-treatment on simvastatin-mediated changes in *APOA1*, *PLTP*, and *GK* expression in Huh7 cells. Gene expression was quantified by qPCR and normalized to the geometric mean of *HPRT* and *B2M*. Data are presented as mean ± SEM, with individual data points shown (n indicated in the figure). Statistical significance is denoted as *p < 0.05, **p < 0.01, ***p < 0.001.

Recent studies have proposed that postprandial absorption of gut microbiota-derived LPS leads to transient elevations in circulating LPS—often termed metabolic endotoxemia (Cani et al., 2007; Mohammad and Thiemermann, 2021). Notably, LPS can be incorporated into intestinally secreted lipoproteins such as chylomicrons, thereby entering systemic circulation (Clemente-Postigo et al., 2012). We measured plasma LPS levels during fasting (ZT03–09) and feeding (ZT15–21) and found that, irrespective of simvastatin treatment, circulating LPS peaked during feeding and was significantly lower during fasting (Figure 4B). To determine whether LPS modulates simvastatin’s effects, we co-treated Huh-7 cells with simvastatin and LPS; this co-treatment abolished simvastatin-induced upregulation of *PLTP*, *APOA1*, and *GK* (Figure 4C). These findings indicate that microbiota-derived LPS, which surges with feeding, may account for the suppression of PPARα activation observed when simvastatin is administered with food (Figures 2A–C).

### LPS-Statin Interaction: A Key Determinant of Oxysterol Metabolism in the Liver

To investigate how feeding-driven LPS might modulate simvastatin’s regulation of PPARα activity—which governs fasting glucose and HDL metabolism—we reanalyzed transcriptomic data from differentiated HepaRG (dHepaRG) cells treated with LPS for six hours (GEO: GSE230325) (Ehle et al., 2024). dHepaRG cells retain key hepatic features, including hepatocyte-like morphology and the expression of metabolic enzymes, nuclear receptors, and drug transporters (Ehle et al., 2024). Our analysis covered both protein-coding and non-coding RNAs, with 43,892 genes detected in total; a volcano plot revealed significant transcriptional changes (Figure 5A).

**Figure 5.**
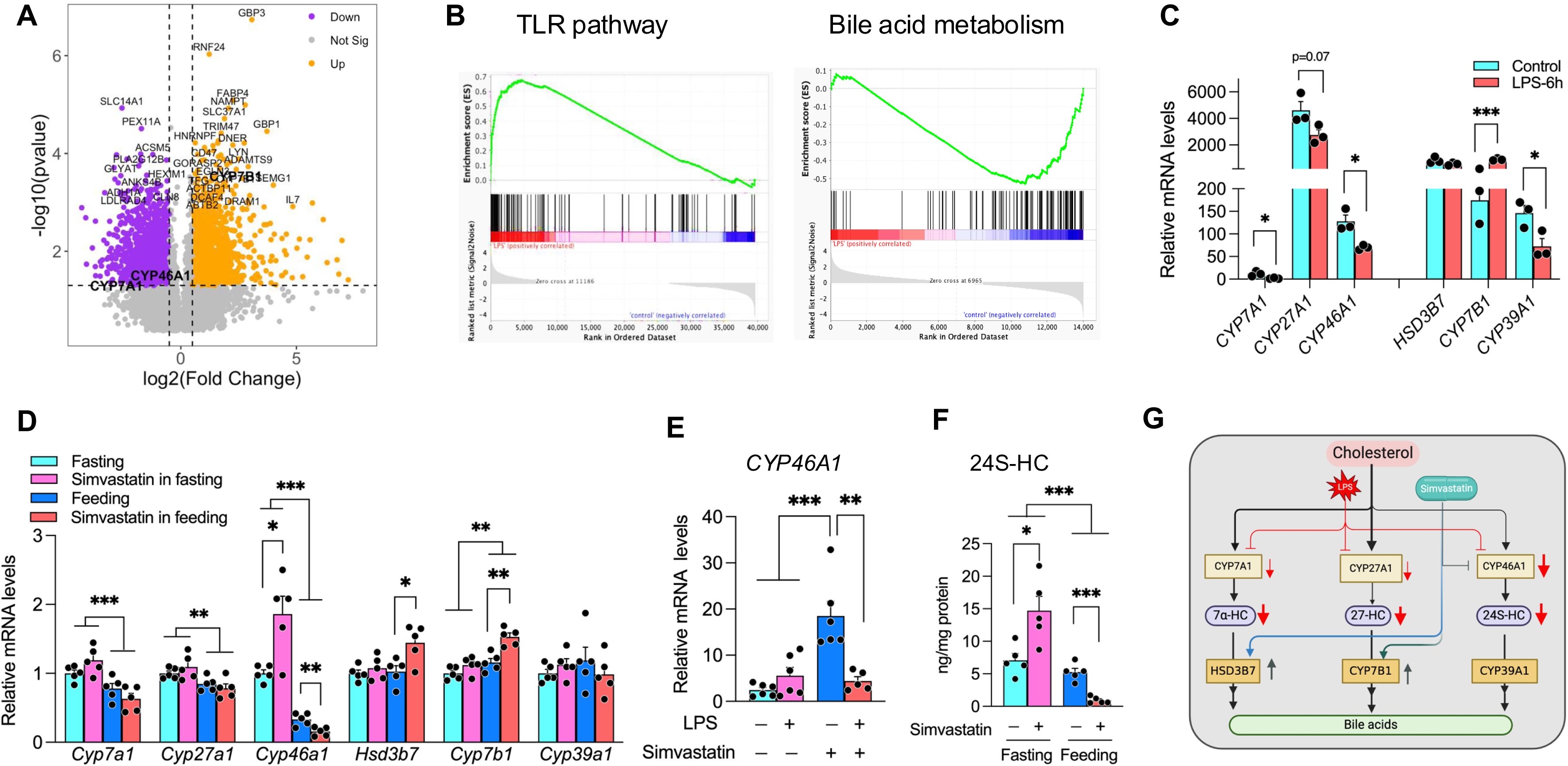
LPS–Statin Interaction Is a Key Determinant of Hepatic Oxysterol Metabolism. **(A)** Volcano plot of differentially expressed genes (both protein-coding and non-coding) in LPS-treated differentiated HepaRG (dHepaRG) cells (GEO: GSE230325). A total of 43,892 genes were detected. Key oxysterol-related enzymes (*CYP7A1*, *CYP46A1*, and *CYP7B1*) are highlighted. The x-axis shows log2 fold change, and the y-axis shows −log10 p-value; dashed lines indicate thresholds for statistical significance (p < 0.05, |log2 fold change| > 1). **(B)** Gene Set Enrichment Analysis (GSEA) of LPS-upregulated genes reveals enrichment in infection-related pathways and Toll-like receptor (TLR) signaling, while genes downregulated by LPS are associated with bile acid metabolism, xenobiotic processing, and fatty acid metabolism. **(C)** Alterations in oxysterol metabolism following LPS treatment in dHepaRG cells. LPS reduces transcript levels of the oxysterol-producing enzymes *CYP7A1*, *CYP27A1*, and *CYP46A1*, while elevating CYP7B1, thus promoting depletion of oxysterols (e.g., 27-HC). **(D)** Feeding-phase simvastatin further modulates bile acid/oxysterol metabolism in mouse liver. qPCR analyses show that feeding decreases oxysterol-producing genes (*CYP7A1*, *CYP27A1*, and *CYP46A1*) and simvastatin coadministration augments oxysterol-catabolizing enzymes (*Cyp7b1*, *Hsd3b7*). **(E)** In Huh7 cells, simvastatin significantly upregulates *CYP46A1*, an effect substantially attenuated by LPS co-treatment. Gene expression values are normalized to *HPRT*. **(F)** Measurement of hepatic 24(S)-hydroxycholesterol (24S-HC) reveals that simvastatin lowers 24S-HC when administered during feeding, whereas it elevates 24S-HC under fasting conditions. **(G)** Schematic model illustrating how feeding-driven LPS signaling synergizes with simvastatin to deplete hepatic oxysterols by suppressing the classic (CYP7A1-mediated) and acidic (*CYP27A1*, *CYP46A1*) pathways and upregulating the oxysterol-metabolizing enzyme *CYP7B1*. Data are presented as mean ± SEM, with individual data points shown (n indicated in the figure). Statistical significance is denoted as *p < 0.05, **p < 0.01, ***p < 0.001.

Gene Set Enrichment Analysis (GSEA) of LPS-upregulated genes showed enrichment in pathways related to infection, cytokine-receptor interactions, and Toll-like receptor signaling— specifically involving the LPS receptor TLR4 (Figure 5B). By contrast, LPS-downregulated genes were enriched in pathways tied to bile acid metabolism, xenobiotic and fatty acid metabolism, peroxisomal functions, and estrogen response (Mootha et al., 2003; Subramanian et al., 2005; Chen et al., 2013; Kuleshov et al., 2016; Xie et al., 2021). We focused on two salient changes: the reduction in bile acid metabolism-related genes and the induction of TLR signaling, aligning with previous reports that LPS suppresses bile acid synthesis and that statins also influence bile acid metabolism (Khovidhunkit et al., 2004; Schonewille et al., 2016). Additionally, LPS-activated TLR4 impacts lipid metabolism, including bile acid pathways (Wang et al., 2024; Castrillo et al., 2003).

In the liver, cholesterol is metabolized into bile acids primarily via two pathways. The classic (neutral) pathway begins with cholesterol 7α-hydroxylase (CYP7A1), which catalyzes the rate-limiting step—generating 7α-hydroxycholesterol (7α-HC)—and proceeds through 3β-hydroxy-Δ5-C27-steroid oxidoreductase (HSD3B7) to form cholic acid and chenodeoxycholic acid (CDCA) (Pandak and Kakiyama, 2019). The acidic pathway predominantly relies on sterol 27-hydroxylase (CYP27A1), which produces (25R)26-hydroxycholesterol (27-HC). This and other oxysterols are further modified by oxysterol 7α-hydroxylase (CYP7B1) to eventually yield CDCA (Pandak and Kakiyama, 2019; Wang et al., 2021). Cholesterol 24(S)-hydroxylase (CYP46A1) generates 24(S)-hydroxycholesterol (24S-HC), which, while predominantly found in neurons, is also expressed in the liver and has regulatory roles in lipid metabolism (Si et al., 2020). Analysis of LPS-treated dHepaRG cells showed a marked decrease in *CYP7A1*, *CYP46A1*, and *CYP39A1*, along with a modest decline in *CYP27A1* (p=0.07). By contrast, *CYP7B1* was upregulated more than fivefold (Figure 5C). These results imply that LPS not only downregulates the classic neutral pathway but also influences the acidic pathway, reducing oxysterol-producing enzymes (*CYP27A1*, *CYP46A1*) while increasing the oxysterol-metabolizing enzyme *CYP7B1*—thereby promoting the depletion of oxysterols such as 27-HC.

We next explored how bile acid and oxysterol metabolism shift during simvastatin treatment in the feeding phase, which induces a surge in LPS (Figure 4B). Relative to fasting, feeding decreased the expression of oxysterol-producing enzymes *Cyp7a1*, *Cyp27a1*, and *Cyp46a1* (Figure 5D). Feeding-phase simvastatin further increased the expression of oxysterol-catabolizing enzymes *Hsd3b7* and *Cyp7b1* (Figure 5D). While feeding-phase simvastatin did not affect *Cyp39a1*, it further suppressed *Cyp46a1*, contrasting with the upregulation observed during fasting (Figure 5D)—indicating that feeding-related factors can reverse simvastatin-induced *Cyp46a1* expression.

Consistently, in Huh-7 cells, simvastatin significantly increased *CYP46A1* expression, an effect that LPS treatment markedly diminished (Figure 5E). In mouse liver, 24S-HC quantification revealed that simvastatin lowered 24S-HC levels during feeding, whereas it increased 24S-HC under fasting conditions (Figure 5F). These results suggest that simvastatin administration with food depletes hepatic oxysterol pools, likely through feeding-driven LPS signals (Figure 5G).

### Feeding-Phase Simvastatin Depletes Oxysterols and Suppresses LXR–SREBP-1c Signaling

Oxysterols—such as 24S-, 25-, and 27-hydroxycholesterol—serve as physiological ligands for liver X receptors (LXRα and LXRβ), activating them at concentrations typically found in vivo (Janowski et al., 1996; Lehmann et al., 1997; Fu et al., 2001). To determine whether feeding-phase simvastatin administration—which depletes hepatic oxysterols—alters LXR activity, we measured the expression of canonical LXR targets. Notably, the genes *Abcg1*, *Abcg5*, and SREBP-1c were significantly repressed when simvastatin was given with food, but not during fasting (Figure 6A). Although activation of TLR3/TLR4 by E. coli or influenza A and ligands like poly I:C (TLR3 ligand) or LPS component lipid A (TLR4 ligand) can block LXR signaling in macrophages via IRF-3 competing for the coactivator p300/CBP, we found that simvastatin co-administration with food significantly downregulated IRF3 and NFκB targets (*Il15*, *Ppp2r1b*, *Crp*, *Saa1*) (Figure 6B). These results suggest that the IRF-3 pathway may not be essential for the synergistic inactivation of LXR by LPS and simvastatin, and that depletion of oxysterols likely represents the key mechanism.

**Figure 6.**
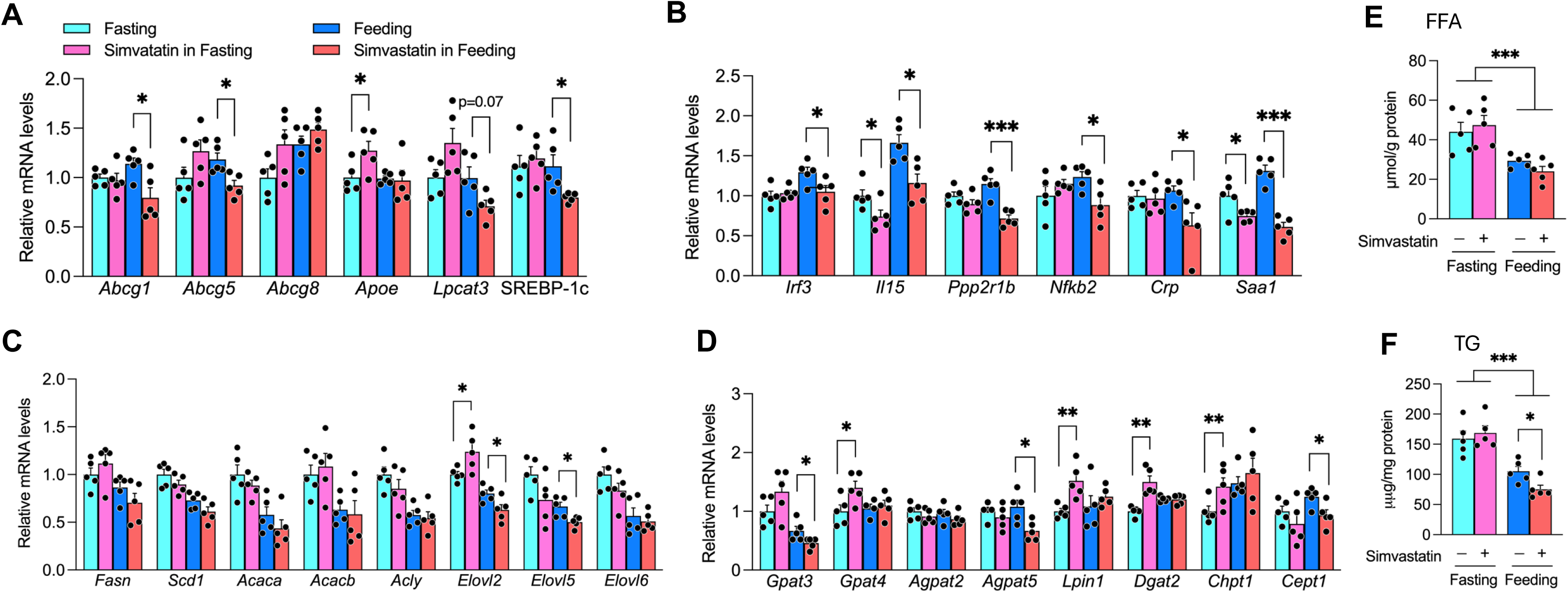
Feeding-Phase Simvastatin Depletes Oxysterols and Suppresses LXR/SREBP-1c Signaling. **(A)** Quantitative PCR analysis of canonical LXR targets in the livers of A/J mice administered simvastatin during feeding or fasting. **(B)** Expression levels of TLR4-dependent NFκB and IRF3 target genes under the same conditions. **(C)** Hepatic expression of SREBP-1c target genes involved in fatty acid biosynthesis. **(D)** Additional SREBP-1c targets that regulate triglyceride and phosphatidylcholine synthesis. **(E, F)** Hepatic free fatty acid (FFA) (E) and triacylglycerol (TG) (F) content in livers of A/J mice treated with simvastatin during feeding versus fasting. Gene expression values are normalized to *Hprt* and presented as mean ± SEM from three independent experiments. Statistical significance is indicated by *p < 0.05, **p < 0.01, ***p < 0.001.

Consistent with prior reports that statin exposure can reduce both membrane-bound and nuclear SREBP-1c in an LXR-dependent manner (DeBose-Boyd et al., 2001; Rong et al., 2017; Liang et al., 2002), we found that simvastatin treatment during feeding—but not fasting—led to decreased SREBP-1c expression (Figure 6A). Reduced SREBP-1c transactivation was evident in the feeding group, where SREBP-1c targets involved in lipogenesis (*Elovl2*, *Elovl5*) and triacylglycerol synthesis (*Gpat3*, *Agpat5*) and phospholipid formation (*Chpt1*, *Cept1*) were suppressed (Figures 6C, D). Quantitatively, simvastatin treatment during feeding did not alter hepatic free fatty acid levels but significantly lowered triacylglycerol content, consistent with diminished SREBP-1c activity (Figures 6E, F).

Because de novo lipogenesis, including phosphatidylcholine synthesis via choline/ethanolamine phosphotransferase 1 (*Cept1*), is required for PPARα activation (Guan et al., 2018; Zayed et al., 2021), these findings support a model in which feeding-driven LPS, together with simvastatin, depletes cellular oxysterol pool and inactivates LXR. This, in turn, hampers SREBP-1c activity and reduces de novo lipogenesis, which is essential for PPARα activation under both postprandial and fasting conditions. The net result is decreased fasting blood glucose and lower HDL cholesterol when simvastatin is co-administered with food.

### TLR4 Deficiency Reverses Simvastatin’s Effects on HDL Cholesterol and Glucose Metabolism in the Feeding State

We observed that LPS suppresses hepatic expression of *CYP7A1*, *CYP27A1*, and *CYP46A1*, thereby priming the liver for further oxysterol depletion by simvastatin during feeding. Previous work has shown that LPS-mediated downregulation of hepatic cytochrome P450 (P450) mRNAs is entirely TLR4-dependent (Chaluvadi et al., 2009). To determine whether TLR4 is also essential for modulating simvastatin’s effects on HDL cholesterol and glucose homeostasis across feeding and fasting states, we used C3H/HeJ mice harboring a spontaneous Tlr4 mutation (Tlr4^Lps-d), which renders them hyporesponsive to LPS (Kamath et al., 2003). These mice, along with wild-type C3H/HeOuJ controls, were fed simvastatin mixed in chow for five weeks. In wild-type C3H/HeOuJ mice, simvastatin co-administration with food significantly improved glucose tolerance while modestly lowering HDL cholesterol (p = 0.08), mirroring the results in A/J mice (Figures 7A, 7B and 1B).

**Figure 7.**
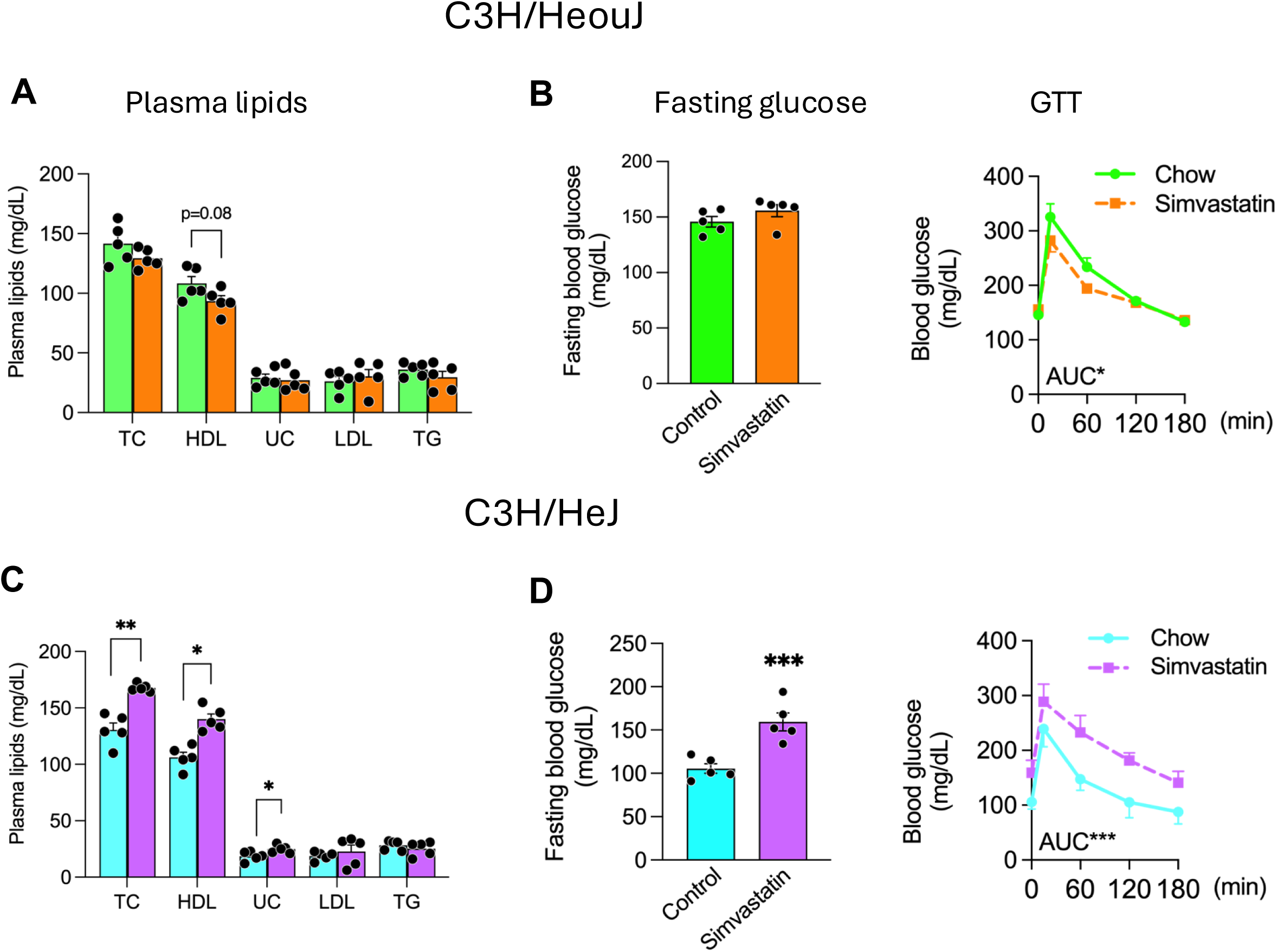
TLR4 Deficiency Alters Simvastatin’s Effects on HDL Cholesterol and Glucose Metabolism. **(A, B)** Plasma lipid and glucose measurements in wild-type C3H/HeOuJ mice co-administered simvastatin with food for 5 weeks. **(C, D)** Plasma lipid and glucose measurements in TLR4-deficient C3H/HeJ mice co-administered simvastatin with food for 5 weeks. Data are presented as mean ± SEM (n = 5). Statistical significance is indicated as *p < 0.05, **p < 0.01, ***p < 0.001.

In contrast, TLR4-deficient C3H/HeJ mice exhibited significantly elevated HDL cholesterol, total cholesterol, and free cholesterol after simvastatin treatment during feeding (Figure 7C). Moreover, the TLR4-mutant mice developed impaired glucose homeostasis under these conditions, as indicated by elevated fasting blood glucose and glucose intolerance (Figure 7D)—a phenotype that closely resembled A/J mice given simvastatin during fasting (Figure 1D). These findings highlight the critical role of TLR4 in mediating feeding-driven LPS modulation of simvastatin’s regulatory effects on HDL cholesterol and glucose metabolism. Without functional TLR4 signaling, the normal synergy between LPS and simvastatin is disrupted, shifting the metabolic outcomes toward elevated HDL and impaired glucose tolerance.

### TLR4 Deficiency Modifies Simvastatin’s Influence on LXR/SREBP-1c/PPARα Pathways

To elucidate how TLR4 modulates simvastatin’s metabolic effects, we compared the expression of key bile acid/oxysterol-metabolizing enzymes in livers from TLR4-deficient (C3H/HeJ) and wild-type (C3H/HeOuJ) mice by quantitative PCR. In wild-type mice, simvastatin treatment suppressed *Cyp46a1* and upregulated *Cyp7b1* (Figure 8A). In contrast, TLR4 deficiency elevated the basal expression of oxysterol-generating enzymes—including *Cyp7a1*, *Cyp27a1*, and *Cyp46a1*. Moreover, in TLR4-deficient livers, simvastatin further increased *Cyp7a1* and *Cyp46a1* expression while downregulating the oxysterol-metabolizing genes *Hsd3b7* and *Cyp39a1* (Figure 8A), suggesting that TLR4 deficiency prevents simvastatin-induced depletion of hepatic oxysterols.

**Figure 8.**
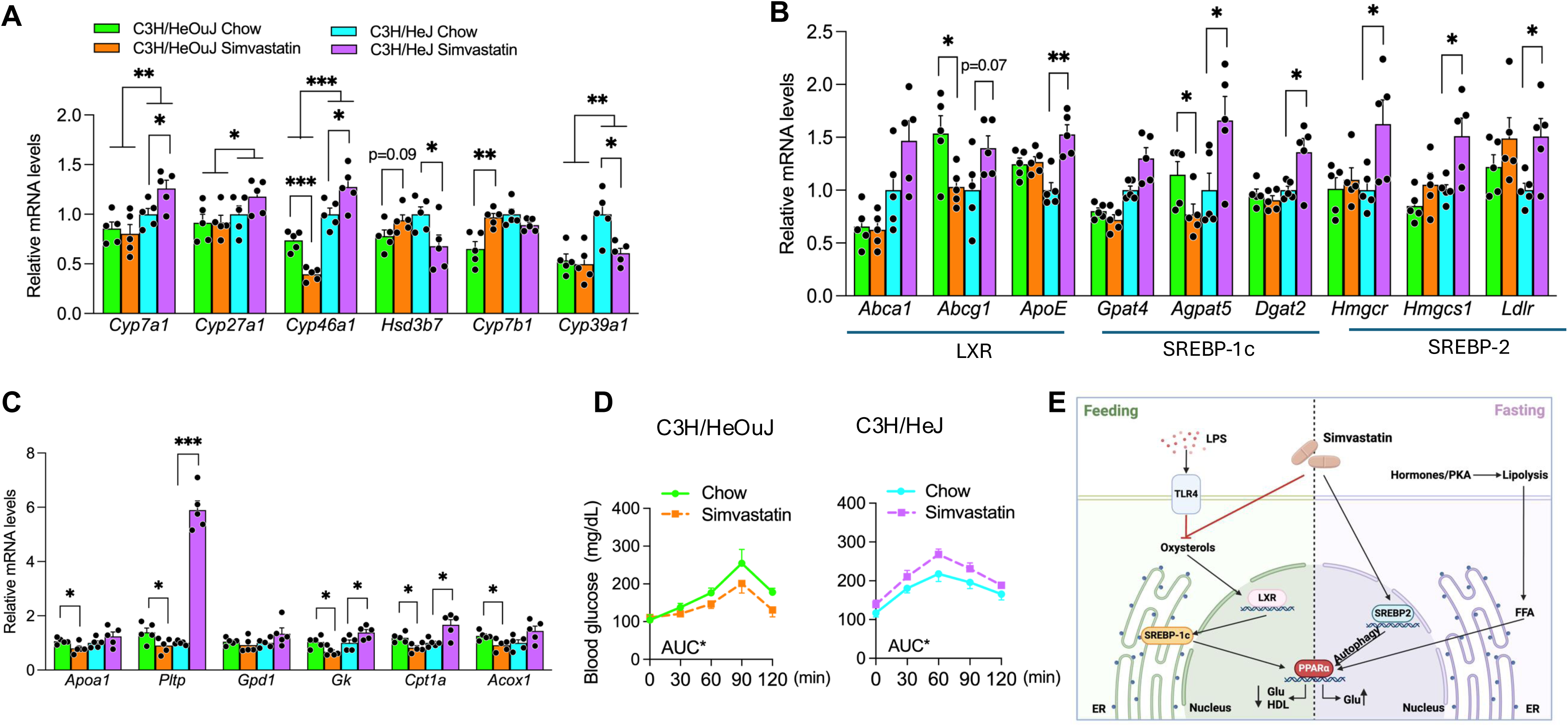
TLR4 Deficiency Modifies Simvastatin’s Influence on LXR/SREBP-1c/PPARα Signaling. **(A)** Quantitative PCR analysis of bile acid/oxysterol metabolism genes in the livers of C3H/HeJ and C3H/HeOuJ mice following simvastatin co-administration with food. **(B)** Expression of LXR, SREBP-1c, and SREBP-2 target genes in the liver of C3H/HeJ and C3H/HeOuJ mice following simvastatin co-administration with food. **(C)** The expression of PPARα-regulated genes involved in HDL metabolism and gluconeogenesis in C3H/HeJ and C3H/HeOuJ mice given simvastatin with food. **(D)** Blood glucose levels measured at 30, 60, 90, and 120 minutes in C3H/HeJ and C3H/HeOuJ mice administered either standard chow or simvastatin-enriched chow. **(E)** Schematic illustration showing how different nutritional states modulate simvastatin’s effects on HDL cholesterol and glucose metabolism via SREBP-2/autophagy/PPARα under fasting and through LPS–statin convergence on the oxysterol/LXR/SREBP-1c/PPARα pathway under feeding. Data are presented as mean ± SEM (n = 5). Statistical significance is denoted as *p < 0.05, **p < 0.01, ***p < 0.001.

TLR4 deficiency also counteracted simvastatin-mediated inhibition of the LXR/SREBP-1c axis. This was evidenced by increased expression of *Apoe*, *Gpat4*, *Cept1*, and *Cpt1a*, which were otherwise reduced in simvastatin-treated wild-type mice (Figure 8B). These findings suggest that, in the absence of TLR4, simvastatin promotes LXR ligand synthesis and activation rather than suppressing it.

Analysis of the PPARα pathway frther showed that simvastatin enhanced PPARα target genes (*Pltp*, *Gk*, *Cpt1a*) in TLR4-deficient mice but downregulated these genes in wild-type controls (Figure 8C). To verify that PPARα contributes to dysglycemia in simvastatin-treated TLR4-deficient mice, we conducted glycerol tolerance tests (Sumara et al., 2012). Glycerol administration led to higher circulating glucose levels in C3H/HeJ mice compared to wild-type mice, confirming that PPARα-driven, glycerol-fueled gluconeogenesis exacerbates glucose intolerance in TLR4-deficient mice given simvastatin (Figure 8D).

Collectively, these results demonstrate that TLR4 is essential for feeding-driven LPS regulation of simvastatin’s metabolic actions. In TLR4-deficient mice, simvastatin’s usual effects on HDL cholesterol and glucose metabolism are reversed due to disrupted SREBP-1c and PPARα pathways, underscoring the pivotal role of TLR4 in modulating simvastatin’s influence on hepatic lipid and glucose homeostasis.

## DISCUSSION

Nutritional states, such as fasting and feeding cycles, profoundly regulate hepatic lipid metabolism largely via SREBP-mediated transcriptional programs that control cholesterol and fatty acid biosynthesis (Stokkan et al., 2001; Vollmers et al., 2009; Greco et al., 2021). Statins are broadly recognized for their capacity to lower LDL-C and reduce cardiovascular risk (Baigent et al., 2010; Fulcher et al., 2015), and their efficacy and side effects are believed to depend on endogenous cholesterol metabolism and its metabolites (Ward et al., 2019). However, the influence of nutritional state on statin-mediated metabolic outcomes remains insufficiently explored. Here, we demonstrate that simvastatin’s effects on HDL-C and glucose metabolism are markedly contingent upon feeding versus fasting conditions. Specifically, simvastatin co-administered with food significantly lowered HDL-C and improved glucose metabolism, whereas simvastatin given during fasting modestly raised HDL-C but substantially impaired glucose tolerance. These data underscore the critical impact of feeding-driven metabolic cues in shaping statin responsiveness.

Statins can elevate HDL-C to varying degrees, although the precise mechanisms remain unclear (Barter et al., 2010). Our results reveal that fasting-versus feeding-dependent differences in HDL-C regulation by simvastatin are mediated by PPARα, a nuclear receptor governing both HDL metabolism and gluconeogenesis. Under fasting conditions, simvastatin robustly induced PPARα and its target genes (e.g., *APOA1* in human hepatocytes and *PLTP* in both human and mouse hepatocytes), which are instrumental in HDL formation (Bouly et al., 2001; Martin et al., 2001; Han et al., 2006). Concurrently, PPARα activation during fasting enhanced expression of genes involved in glycerol-fueled gluconeogenesis (*Gpd1*, *Gk*, and *Aqp9*), explaining the observed glucose intolerance. Conversely, co-administration of simvastatin with food suppressed PPARα activity, downregulating key targets involved in HDL production and gluconeogenesis. These findings indicate that PPARα acts as a metabolic switch, integrating feeding signals to modulate the lipid and glycemic responses induced by statins.

During fasting, FFAs mobilized from adipose tissue provide substrates and ligands for hepatic PPARα (Bideyan et al., 2021). Although synthetic PPARα ligands (e.g., fibrates) strongly induce HDL-related genes, long-chain FFAs often have minimal or even inhibitory effects (Kuang et al., 2012). In our study, oleic acid alone was insufficient to induce *APOA1* expression; however, when combined with simvastatin, oleic acid synergistically activated PPARα, markedly inducing *APOA1*. Mechanistic analyses suggest that simvastatin enhances PPARα activity via SREBP-2-dependent autophagy, potentially promoting the degradation of nuclear receptor corepressors such as NCoR1 and HDAC3 (Saito et al., 2019; Iershov et al., 2019). These results imply that, during fasting, statins may potentiate PPARα-driven metabolism by stimulating autophagy to increase HDL and gluconeogenesis (Figure 8E).

Our data reveal that simvastatin co-administered with food exerts effects opposite to those observed under fasting, including a reduction in HDL-C and improved glucose tolerance. We identify feeding-induced LPS as a key mediator of this divergence. Dietary intake triggers a transient surge in circulating LPS (Cani et al., 2007), which subsides during fasting. We propose that LPS, in conjunction with simvastatin, depletes hepatic oxysterol pools by altering the expression of enzymes responsible for oxysterol biosynthesis and metabolism. The resulting decrease in LXR activity suppresses SREBP-1c and de novo lipogenesis, both of which are integral to PPARα activation in postprandial and fasting states (Figure 8E) (Guan et al., 2018; Zayed et al., 2021).

Feeding-driven LPS also disrupts bile acid metabolism by downregulating *CYP7A1*, *CYP27A1*, and *CYP46A1*, reducing oxysterol formation. Simvastatin further amplifies this effect by upregulating oxysterol-metabolizing enzymes such as *Cyp7b1*, leading to oxysterol depletion and sustained LXR inactivation. Previous findings on atorvastatin-induced *Cyp7a1* upregulation involved doses and half-lives distinct from those of simvastatin (McIver and Siddique, 2020; Talreja et al., 2023; Fu et al., 2014; Schonewille et al., 2016), potentially explaining discrepancies in the literature. These observations underscore a novel mechanism in which fluctuations in postprandial endotoxemia modulate SREBP-1c-mediated lipogenesis and statin-induced changes in lipid and glucose metabolism. This aligns with evidence that antibiotic-driven depletion of gut microbiota attenuates statin-induced reductions in HDL-C, highlighting gut microbiota-derived LPS as a critical determinant of statin action (Zimmermann et al., 2020). Our work further implicates TLR4 as a critical mediator of simvastatin’s metabolic effects during feeding. To elucidate its role, TLR4-deficient C3H/HeJ mice (harboring a loss-of-function mutation) were compared with wild-type C3H/HeOuJ controls. In wild-type mice, simvastatin administered with food improved glucose tolerance and modestly reduced HDL-C. By contrast, under the same conditions, TLR4-deficient mice exhibited elevated total and HDL-C along with impaired glucose homeostasis, indicating that intact TLR4 signaling is essential for the typical metabolic actions of simvastatin during feeding. At the molecular level, TLR4 deficiency prevented the feeding-driven LPS suppression of oxysterol biosynthesis, thereby maintaining hepatic expression of *Cyp7a1*, *Cyp27a1*, and *Cyp46a1*. This preservation of oxysterols sustained LXR activity, counteracting simvastatin’s inhibitory effect on Srebf1 and its downstream lipogenic genes. Moreover, TLR4 deficiency rescued PPARα activation and its targets, including *Pltp*, *Gk*, and *Cpt1a*, underscoring the intricate interplay between LPS signaling, statin metabolism, and nuclear receptor regulation.

An intriguing finding was the divergence in glycemic regulation between wild-type and TLR4-deficient mice. Whereas simvastatin co-administration with food enhanced glucose tolerance in wild-type mice, it worsened glucose intolerance in TLR4-deficient mice, mirroring effects seen with fasting. Glycerol tolerance tests revealed that TLR4 deficiency amplified simvastatin-induced PPARα activation, fueling glycerol-driven gluconeogenesis and hyperglycemia. These findings pinpoint TLR4 as a crucial node linking LPS, statin action, and systemic metabolic homeostasis.

In summary, our study illustrates how feeding and fasting states critically shape simvastatin’s impact on HDL and glucose metabolism via distinct molecular pathways. Under fasting conditions, simvastatin promotes PPARα activation potentially related to autophagy-driven degradation of nuclear receptor corepressors, increasing HDL-C but also elevating gluconeogenesis. In contrast, feeding-triggered LPS-TLR4 signaling dampens PPARα activity, reducing HDL-C while enhancing glucose tolerance (Figure 8E). These insights offer a mechanistic framework for the interindividual variability in statin responses and underscore the importance of timing statin administration with feeding behaviors to optimize efficacy and minimize metabolic side effects.

## Materials and Methods

### Animals

Male A/J, C3H/HeJ, and C3H/HeOuJ mice were obtained from The Jackson Laboratory (Bar Harbor, ME) and housed at ambient temperature on a 12-hour light/dark cycle. Animal experiments were conducted following protocols approved by the University of Maryland Baltimore Institutional Animal Care and Use Committee.

### Simvastatin Administration

To investigate the differential effects of simvastatin during fasting and feeding states, 8–10-week-old mice were maintained on standard chow. Mice typically consumed ∼80% of their food during the dark (active) phase (ZT12–ZT24) and exhibited minimal intake during the light (rest) phase (ZT0–ZT12) (Turek et al., 2005; Brown et al., 2021). Two treatment protocols were employed: dietary administration and oral gavage.

In the dietary administration protocol, mice were provided either a control chow diet (Diet D1001; 10 kcal% fat, 20 kcal% protein, 70 kcal% carbohydrate; Research Diets, New Brunswick, NJ) or chow supplemented with simvastatin (0.1 g/kg food weight; Formulation D11060903i, Research Diets). This dosage approximates an 80 mg/day human equivalent (Zhang et al., 2023). In the oral gavage protocol, mice received simvastatin at 16 mg/kg body weight, suspended in 1% methylcellulose in 1× PBS, administered once daily at 10:00 AM (ZT04) using 20-gauge plastic feeding tubes (Lastuvkova et al., 2021). The timing coincided with the light-phase fasting period to compare simvastatin effects under fasting versus feeding conditions.

Mice were housed on a 12-hour light/dark cycle (lights on at 6:00 AM [ZT0] and off at 6:00 PM [ZT12]) with ad libitum access to food and water. Plasma and tissue samples were collected after five weeks of treatment to capture metabolic states at distinct time points: ZT09 (3:00 PM), five hours post-gavage for the fasting-phase group, and ZT05 (11:00 AM), five hours post-feeding for the dietary administration group.

### Glucose Tolerance Test (GTT)

Mice in the fasting-phase group were fasted for five hours following the last oral gavage and underwent GTT at ZT09 (3:00 PM). Mice in the feeding-phase group were fasted for five hours post-feeding, with GTT performed at ZT05 (11:00 AM). Baseline fasting glucose levels were measured from blood samples before administering glucose. Mice were injected intraperitoneally with glucose (2 g/kg body weight), and blood glucose levels were measured via tail nick at 15, 60, 120, and 180 minutes post-injection using an AlphaTRAK glucometer (Zoetis, Parsippany, NJ). The area under the curve (AUC) for glucose tolerance was determined using trapezoidal integration (Zhang et al., 2024; Solberg et al., 2006).

### Glycerol Tolerance Measurement

Mice were fasted for five hours and administered glycerol (1.5 g/kg body weight) intraperitoneally. Plasma glucose levels were measured at specified time points to evaluate glycerol tolerance (Kim et al., 2017).

### Cell Culture

Huh7 human hepatoma cells, provided by Dr. Liqing Yu (Yang et al., 2020), were maintained at 37 °C in a humidified atmosphere containing 5% CO₂. Cells were cultured in Dulbecco’s Modified Eagle Medium (DMEM) supplemented with 10% fetal bovine serum (FBS), 100 U/ml penicillin, and 100 µg/ml streptomycin until subconfluent.

For fasting mimetic conditions, cells were washed twice with 1× PBS and incubated overnight in serum-free DMEM. Cells were then treated with either 50 µM oleic acid (Millipore Sigma, C18:1; #O1383), 500 µM leucine, or 10 nM insulin, with or without 5 µM simvastatin, for 24 hours. Total RNA was extracted using Trizol according to the manufacturer’s instructions. In serum-replete conditions, cells were cultured in DMEM with 10% FBS and treated with either 50 µM oleic acid, 250 µM leucine, or 10 nM insulin, with or without 10 µM simvastatin, for 8 hours. RNA extraction followed the same procedure.

### Biochemical Measurements

Plasma cholesterol fractions (HDL-C, total cholesterol [TC], LDL-C, unesterified cholesterol [UC]) and triglyceride levels were quantified using enzymatic kits (Millipore Sigma) according to the manufacturer’s protocols.

### RNA extraction and quantitative real-time PCR

Liver tissues snap-frozen in liquid nitrogen upon dissection and further stored at−80 °C were cut to ∼50 mg piece on dry ice, and homogenized with Tissue Homogenizer in TRIzol by Invitrogen. RNA was extracted and purified according to manufacturer’s specifications, and 1–3 µg were taken for cDNA synthesis (iScript™ Advanced cDNA Synthesis Kit, Biorad). The quantitative PCR was carried out using iTaq™ Universal SYBR® Green Supermix (Biorad) using a StepOnePlus instrument (Applied Biosystems) or with CFX96 Real-Time-PCR detection system (Bio-Rad), and gene expression was normalized to Actb expression. The list of primers used for real-time PCR is given in Table 1. Expression levels were normalized to the housekeeping genes *B2M* and *HPRT* for both human Huh7 cells and mouse liver samples.

**Table 1.**
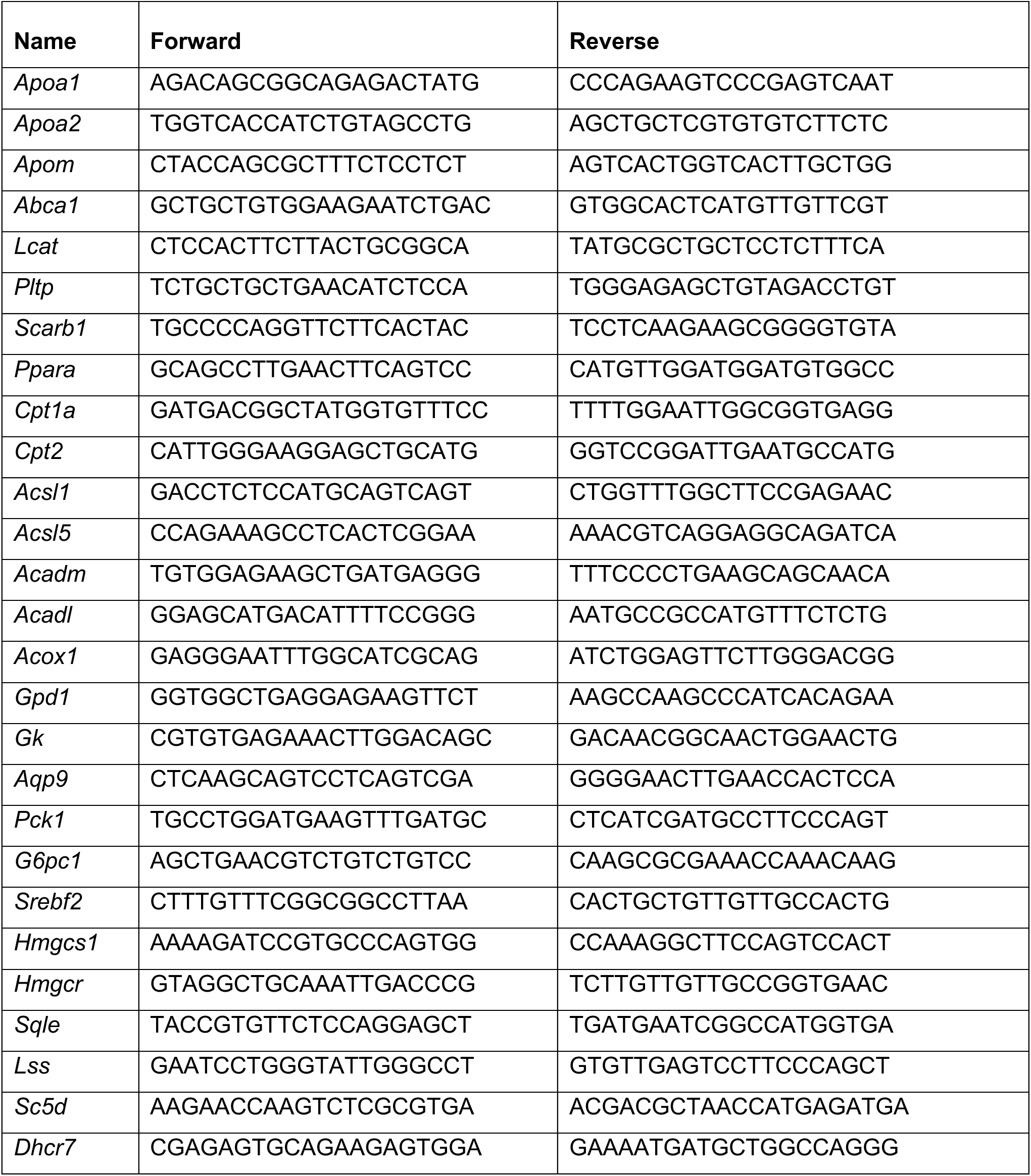
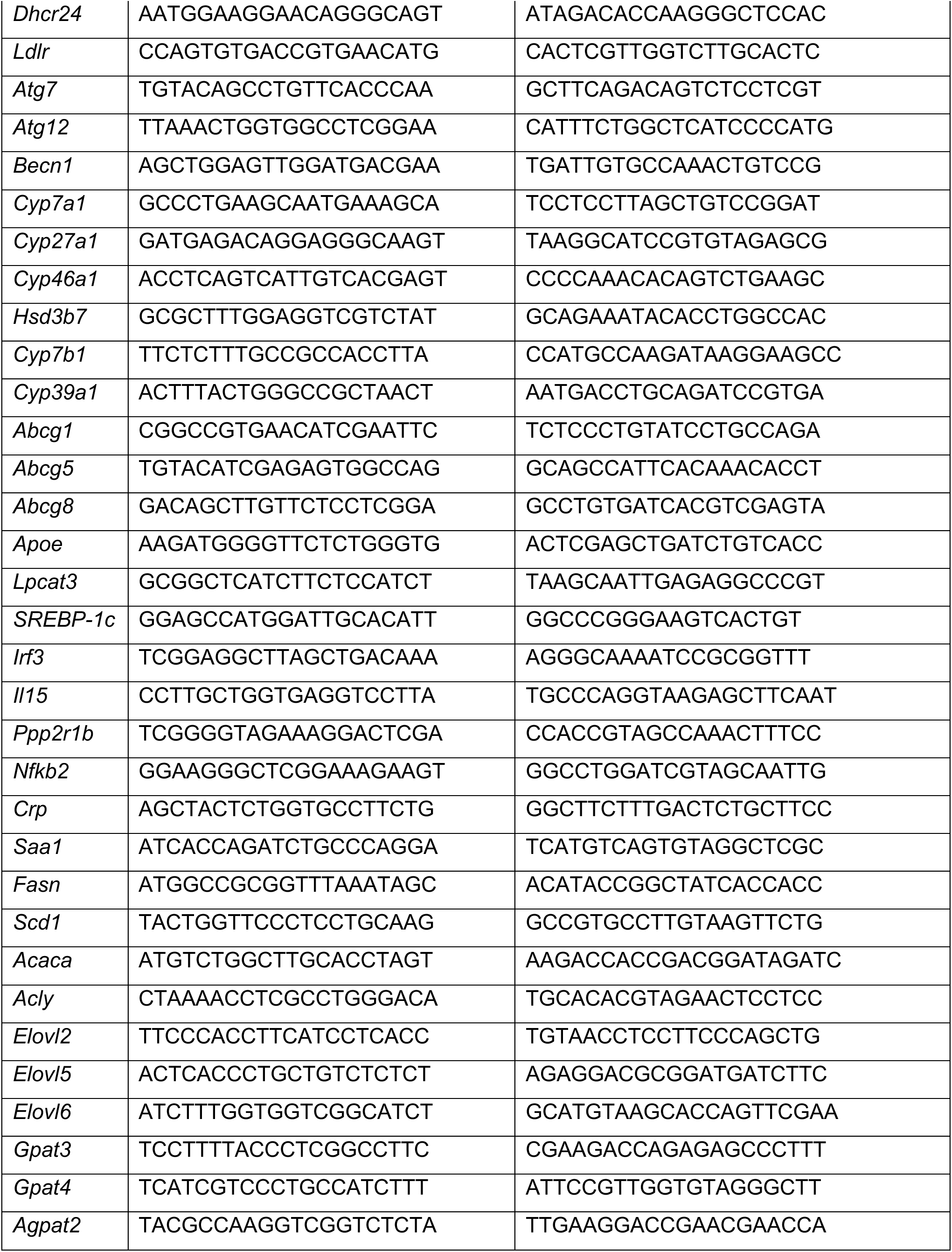
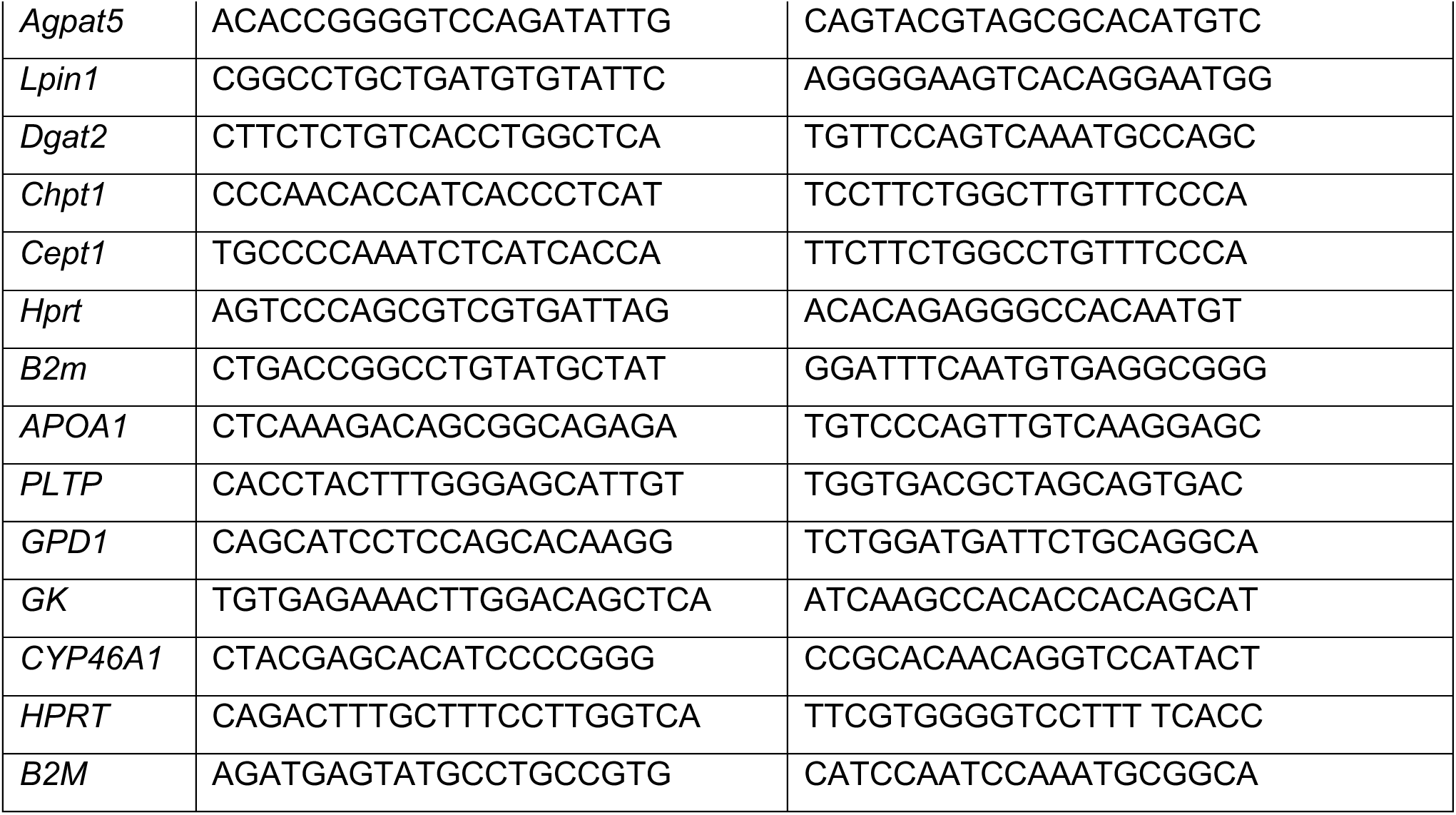
Primers for Quantitative PCR.

### Hepatic Lipid Analysis

Liver lipids were extracted using a modification of the Bligh and Dyer method (Zhang et al., 2019). Approximately 100 mg of liver tissue was homogenized in 4 ml of chloroform/methanol (2:1) and incubated overnight at room temperature. Following the addition of 800 µl of 0.9% saline, the mixture was centrifuged at 2,000 g for 10 minutes. The organic phase was collected, dried under vacuum, and dissolved in butanol/(Triton X-100/methanol [2:1]) (30:20). Triglyceride content was determined using a colorimetric assay (Sigma), and free fatty acids were quantified using FFA kits (Wako Chemicals).

### 24(S)-Hydroxycholesterol Quantification

Hepatic 24(S)-hydroxycholesterol (24S-HC) levels were measured using a 24(S)-hydroxycholesterol ELISA kit (Abcam, USA, #ab204530). Approximately 200 mg of liver tissue was homogenized in 3 mL of 95% ethanol. The homogenate was centrifuged, and the resulting pellets were resuspended in 1 mL of ethanol/dichloromethane (1:1, v/v). The mixture was centrifuged again, and the supernatants from both centrifugation steps were combined. The pooled supernatants were evaporated to dryness using a rotary evaporator. The dried samples were rehydrated with 16 μL of 95% ethanol and 484 μL of assay buffer at room temperature. After appropriate dilution in assay buffer, 24S-HC levels were quantified following the manufacturer’s protocol.

### Statistical Analysis

Statistical analyses were performed using Prism (GraphPad Software, San Diego, CA). Two-way ANOVA was applied to experiments involving multiple groups. Significant interactions (p < 0.05) were followed by pairwise Student’s t-tests. Results of pairwise comparisons are presented in figures, with asterisks denoting levels of significance. Error bars indicate the standard error of the mean (SEM).

## REFERENCES

Alakhali, K.M., Alzubaidi, M.A. and Al Mekhlafi, F.S. (2018) Effect of timing of statin administration on lipid profile: a systematic review and meta analysis. J. Clin. Lipidol. 12, 816–824.

Awad, K., Serban, M.C., Penson, P., Mikhailidis, D.P., Toth, P.P., Jones, S.R., Borow, K.M., Blaha, M.J., Lip, G.Y.H., Banach, M., et al. (2017) Effects of morning vs. evening statin administration on lipid profile: a systematic review and meta analysis. J. Clin. Lipidol. 11, 972–985.e9.

Baigent, C., Blackwell, L., Emberson, J., Holland, L.E., Reith, C., Bhala, N., Peto, R., Barnes, E.H., Keech, A. and Simes, J., et al. (2010) Efficacy and safety of more intensive lowering of LDL cholesterol: a meta analysis of data from 170,000 participants in 26 randomised trials. Lancet 376, 1670–1681.

Bideyan, L., Nagari, R. and Tontonoz, P. (2021) Hepatic transcriptional responses to fasting and feeding. Genes Dev. 35, 635–657.

Bouly, M., Masson, D., Gross, B., Jiang, X.C., Fievet, C., Castro, G., Tall, A.R., Fruchart, J.C., Staels, B., Lagrost, L. and Luc, G. (2001) Induction of the phospholipid transfer protein gene accounts for the high density lipoprotein enlargement in mice treated with fenofibrate. J. Biol. Chem. 276, 25841–25847.

Brown, S.A., et al. (2021) Circadian rhythms in the mouse liver: from gene expression to metabolism and disease. J. Hepatol. 75, 279–291.

Cani, P.D., Amar, J., Iglesias, M.A., Poggi, M., Knauf, C., Bastelica, D., Neyrinck, A.M., Fava, F., Tuohy, K.M. and Chabo, C., et al. (2007) Metabolic endotoxemia initiates obesity and insulin resistance. Diabetes 56, 1761–1772.

Castrillo, A., Joseph, S.B., Marathe, C., Mangelsdorf, D.J. and Tontonoz, P. (2003) Crosstalk between LXR and Toll like receptor signaling mediates bacterial and viral antagonism of cholesterol metabolism. Mol. Cell 12, 805–816.

Chakravarthy, M.V., Lodhi, I.J., Yin, L., Malapaka, R.R., Xu, H.E., Turk, J. and Semenkovich, C.F. (2009) Identification of a physiologically relevant endogenous ligand for PPARŒ± in liver. Cell 138, 476–488.

Chaluvadi, M.R., Nyagode, B.A., Kinloch, R.D. and Morgan, E.T. (2009) TLR4 dependent and independent regulation of hepatic cytochrome P450 in mice with chemically induced inflammatory bowel disease. Biochem. Pharmacol. 77, 464–471.

Chen, E.Y., Tan, C.M., Kou, Y., Duan, Q., Wang, Z., Meirelles, G.V., Clark, N.R. and Ma’ayan, A. (2013) Enrichr: interactive and collaborative HTML5 gene list enrichment analysis tool. BMC Bioinformatics 14, 128.

Chen, W., Chen, G., Head, D.L., Mangelsdorf, D.J. and Russell, D.W. (2007) Enzymatic reduction of oxysterols impairs LXR signaling in cultured cells and the livers of mice. Cell Metab. 5, 73–79.

Cholesterol Treatment Trialists’ (CTT) Collaboration. (2024) Effects of statin therapy on diagnoses of new onset diabetes and worsening glycaemia in large scale randomised blinded statin trials: an individual participant data meta analysis. Lancet Diabetes Endocrinol. 12, 306–319.

Clemente Postigo, M., Queipo Ortuño, M.I., Murri, M., Boto Ordoñez, M., Perez Martinez, P., Andres Lacueva, C., Cardona, F. and Tinahones, F.J. (2012) Endotoxin increase after fat overload is related to postprandial hypertriglyceridemia in morbidly obese patients. J. Lipid Res. 53, 973–978.

DeBose Boyd, R.A., Ou, J., Goldstein, J.L. and Brown, M.S. (2001) Expression of sterol regulatory element binding protein 1c (SREBP 1c) mRNA in rat hepatoma cells requires endogenous LXR ligands. Proc. Natl. Acad. Sci. U.S.A. 98, 1477–1482.

Ehle, C., Iyer Bierhoff, A., Wu, Y., Xing, S., Kiehntopf, M., Mosig, A.S., Godmann, M. and Heinzel, T. (2024) Downregulation of HNF4A enables transcriptomic reprogramming during the hepatic acute phase response. Commun. Biol. 7, 589.

Fu, X., Menke, J.G., Chen, Y., Zhou, G., MacNaul, K.L., Wright, S.D., Sparrow, C.P. and Lund, E.G. (2001) 27 hydroxycholesterol is an endogenous ligand for liver X receptor in cholesterol loaded cells. J. Biol. Chem. 276, 38378–38387.

Fulcher, J., O’Connell, R., Voysey, M., Emberson, J., Blackwell, L., Mihaylova, B., Simes, J., Collins, R., Kirby, A., Colhoun, H., et al. (2015) Efficacy and safety of LDL lowering therapy among men and women: meta analysis of individual data from 174,000 participants in 27 randomised trials. Lancet 385, 1397–1405.

Greco, C.M., Koronowski, K.B., Smith, J.G., Shi, J., Kunderfranco, P., Carriero, R., Chen, S., Samad, M., Sacchi, S. and Sassone Corsi, P. (2021) Integration of feeding behavior by the liver circadian clock reveals network dependency of metabolic rhythms. Sci. Adv. 7, eabe0876.

Guan, D., Xiong, Y., Borck, P.C., Jang, C., Doulias, P.T., Papazyan, R., Fang, B., Jiang, C., Zhang, Y. and Briggs, E.R., et al. (2018) Diet induced circadian enhancer remodeling synchronizes opposing hepatic lipid metabolic processes. Cell 174, 831–842.e12.

Guo, Q., Zhao, Y., Li, J., Liu, J., Yang, X., Guo, X., Kuang, M., Xia, H., Zhang, Z. and Cao, L., et al. (2021) Induction of alarmin S100A8/A9 mediates activation of aberrant neutrophils in the pathogenesis of COVID 19. Cell Host Microbe 29, 222–235.e4.

Han, C.Y., Chiba, T., Campbell, J.S., Fausto, N., Chaisson, M., Orasanu, G., Plutzky, J. and Chait, A. (2006) Reciprocal and coordinate regulation of serum amyloid A versus apolipoprotein A I and paraoxonase 1 by inflammation in murine hepatocytes. Arterioscler. Thromb. Vasc. Biol. 26, 1806–1813.

Horton, J.D., Goldstein, J.L. and Brown, M.S. (1998) SREBPs: activators of the complete program of cholesterol and fatty acid synthesis in the liver. Genes Dev. 12, 2831–2843.

Horton, J.D., Goldstein, J.L. and Brown, M.S. (1998) SREBPs: activators of the complete program of cholesterol and fatty acid synthesis in the liver. J. Clin. Invest. 101, 2331–2339.

Iershov, A., Nemazanyy, I., Alkhoury, C., Girard, M., Barth, E., Cagnard, N., Montagner, A., Chretien, D., Rugarli, E.I. and Guillou, H., et al. (2019) The class 3 PI3K coordinates autophagy and mitochondrial lipid catabolism by controlling nuclear receptor PPARŒ±. Nat. Commun. 10, 1566.

Izquierdo Palomares, J.M., Gracia Marco, T. and Aguilar, F. (2016) Statin timing and lipid profile: systematic review and meta analysis. Cochrane Database Syst. Rev. 2016, CD011231.

Janowski, B.A., Willy, P.J., Devi, T.R., Falck, J.R. and Mangelsdorf, D.J. (1996) An oxysterol signalling pathway mediated by the nuclear receptor LXRŒ±. Nature 383, 728–731.

Kamath, A.B., Alt, J., Debbabi, H. and Behar, S.M. (2003) Toll like receptor 4 defective C3H/HeJ mice are not more susceptible than other C3H substrains to infection with Mycobacterium tuberculosis. Infect. Immun. 71, 4112–4118.

Kim, M., Astapova, I.I., Flier, S.N., Hannou, S.A., Doridot, L., Sargsyan, A., Kou, H.H., Fowler, A.J., Liang, G. and Herman, M.A. (2017) Intestinal, but not hepatic, ChREBP is required for fructose tolerance. JCI Insight 2(24), e96703.

Kim, S., Lee, S., Lee, H. and Lee, E. (2013) Effect of statin administration time on the pharmacokinetics and pharmacodynamics of HMG CoA reductase inhibitors: a systematic review. Clin. Ther. 35, 1786–1795.e5.

Khovidhunkit, W., Kim, M.S., Memon, R.A., Shigenaga, J.K., Moser, A.H., Feingold, K.R. and Grunfeld, C. (2004) Effects of infection and inflammation on lipid and lipoprotein metabolism: mechanisms and consequences to the host. J. Lipid Res. 45, 1169–1196.

Kopito, L., Barrows, C.H. and Van Bruggen, J.T. (1982) Diurnal variations in hepatic HMG CoA reductase activity and plasma mevalonate in rats. Am. J. Physiol. 242, E136–E139.

Kuleshov, M.V., Jones, M.R., Rouillard, A.D., Fernandez, N.F., Duan, Q., Wang, Z., Koplev, S., Jenkins, S.L., Jagodnik, K.M. and Lachmann, A., et al. (2016) Enrichr: a comprehensive gene set enrichment analysis web server 2016 update. Nucleic Acids Res. 44, W90–W97.

Kuo, A. and Hla, T. (2024) Regulation of cellular and systemic sphingolipid homeostasis. Nat. Rev. Mol. Cell Biol. 25, 802–821.

Lehmann, J.M., Kliewer, S.A., Moore, L.B., Smith Oliver, T.A., Oliver, B.B., Su, J.L., Sundseth, S.S., Winegar, D.A., Blanchard, D.E. and Spencer, T.A., et al. (1997) Activation of the nuclear receptor LXR by oxysterols defines a new hormone response pathway. J. Biol. Chem. 272, 3137–3140.

Lastuvkova, H., Faradonbeh, F.A., Schreiberova, J., Hroch, M., Mokry, J., Faistova, H., Nova, Z., Hyspler, R., Igreja Sa, I.C., Nachtigal, P., Stefela, A., Pavek, P. and Micuda, S. (2021) Atorvastatin modulates bile acid homeostasis in mice with diet induced nonalcoholic steatohepatitis. Int. J. Mol. Sci. 22(12), 6468.

Liang, G., Yang, J., Horton, J.D., Hammer, R.E., Goldstein, J.L. and Brown, M.S. (2002) Diminished hepatic response to fasting/refeeding and liver X receptor agonists in mice with selective deficiency of sterol regulatory element binding protein 1c. J. Biol. Chem. 277, 9520–9528.

Liu, Y., Wang, Z., Jin, H., Cui, L., Huo, B., Xie, C., Li, J., Ding, H., Zhang, H. and Xiong, W., et al. (2024) Squalene epoxidase catalyzed 24(S),25 epoxycholesterol synthesis promotes trained immunity mediated antitumor activity. Cell Rep. 43, 114094.

Maqsood, M.H., Jamshaid, T. and Javed, A. (2022) Effect of morning versus evening dosing of statins on lipid profile: a systematic review and meta analysis. Cureus 14, e21345.

Martin, G., Duez, H., Blanquart, C., Berezowski, V., Poulain, P., Fruchart, J.C., Najib Fruchart, J., Glineur, C. and Staels, B. (2001) Statin induced inhibition of the Rho signaling pathway activates PPARŒ± and induces HDL apoA I. J. Clin. Invest. 107, 1423–1432.

Martin, P.D., Mitchell, P.D., Schneck, D.W. and Warwick, M.J. (2002) Pharmacodynamic effects and pharmacokinetics of a new HMG CoA reductase inhibitor, rosuvastatin, after morning or evening administration in healthy volunteers. Br. J. Clin. Pharmacol. 54, 472–477.

McLean, A.J., Le Couteur, D.G. and Hilmer, S.N. (2018) The impact of dosing time on the efficacy and safety of statins: a systematic review and meta analysis. Br. J. Clin. Pharmacol. 84, 2458–2473.

McNamara, D.J., Kolb, R., Parker, T.S., Batwin, H., Samuel, P., Brown, C.D. and Ahrens, E.H. Jr. (1985) HMG CoA reductase activity and plasma mevalonate as predictors of plasma response to cholesterol lowering diets. J. Lipid Res. 26, 1034–1041.

Mootha, V.K., Lindgren, C.M., Eriksson, K.F., Subramanian, A., Sihag, S., Lehar, J., Puigserver, P., Carlsson, E., Ridderstråle, M. and Laurila, E., et al. (2003) PGC 1alpha responsive genes involved in oxidative phosphorylation are coordinately downregulated in human diabetes. Nat. Genet. 34, 267–273.

Mohammad, S. and Thiemermann, C. (2021) Role of metabolic endotoxemia in systemic inflammation and potential interventions. Front. Immunol. 11, 594150.

Nakamoto, I., Uiji, S., Okata, R., Endo, H., Tohyama, S., Nitta, R., Hashimoto, S., Matsushima, Y., Wakimoto, J. and Hashimoto, S., et al. (2021) Diurnal rhythms of urine volume and electrolyte excretion in healthy young men under differing intensities of daytime light exposure. Sci. Rep. 11, 13097.

Parker, T.S., McNamara, D.J., Brown, C., Garrigan, O., Kolb, R. and Batwin, H. (1984) Plasma mevalonate as a measure of cholesterol synthesis in man. J. Clin. Invest. 74, 795–804.

Pang, S., Tang, H., Zhuo, S., Zang, Y.Q. and Le, Y. (2010) Regulation of fasting fuel metabolism by Toll like receptor 4. Diabetes 59, 3041–3048.

Patel, S.J., Liu, N., Piaker, S., Gulko, A., Andrade, M.L., Heyward, F.D., Sermersheim, T., Edinger, N., Srinivasan, H. and Emont, M.P., et al. (2022) Hepatic IRF3 fuels dysglycemia in obesity through direct regulation of Ppp2r1b. Sci. Transl. Med. 14, eabh3831.

Patsouris, D., Mandard, S., Voshol, P.J., Escher, P., Tan, N.S., Havekes, L.M., Koenig, W., März, W., Tafuri, S., Wahli, W., Müller, M. and Kersten, S. (2004) PPARalpha governs glycerol metabolism. J. Clin. Invest. 114, 94–103.

Popják, G., Boehm, G. and Parker, T.S. (1981) Effect of feeding and fasting on the diurnal rhythm of hepatic HMG CoA reductase activity in the rat. J. Lipid Res. 22, 753–761.

Prasad, S.S., Garg, A. and Agarwal, A.K. (2011) Enzymatic activities of the human AGPAT isoform 3 and isoform 5: localization of AGPAT5 to mitochondria. J. Lipid Res. 52, 451–462.

Repa, J.J., Liang, G., Ou, J., Bashmakov, Y., Lobaccaro, J.M., Shimomura, I., Shan, B., Brown, M.S., Goldstein, J.L. and Mangelsdorf, D.J. (2000) Regulation of mouse sterol regulatory element binding protein 1c gene (SREBP 1c) by oxysterol receptors, LXRalpha and LXRbeta. Genes Dev. 14, 2819–2830.

Rong, S., Cortés, V.A., Rashid, S., Anderson, N.N., McDonald, J.G., Liang, G., Moon, Y.A. and Hammer, R.E. (2017) Expression of SREBP 1c requires SREBP 2 mediated generation of a sterol ligand for LXR in livers of mice. eLife 6, e25015.

Saito, T., Kuma, A., Sugiura, Y., Ichimura, Y., Obata, M., Kitamura, H., Okuda, S., Lee, H.C., Ikeda, K. and Kanegae, Y., et al. (2019) Autophagy regulates lipid metabolism through selective turnover of NCoR1. Nat. Commun. 10, 1567.

Seo, Y.K., Jeon, T.I., Chong, H.K., Biesinger, J., Xie, X. and Osborne, T.F. (2011) Genome wide localization of SREBP 2 in hepatic chromatin predicts a role in autophagy. Cell Metab. 13, 367–375.

Shi, H., Kokoeva, M.V., Inouye, K., Tzameli, I., Yin, H. and Flier, J.S. (2006) TLR4 links innate immunity and fatty acid-induced insulin resistance. J. Clin. Invest. 116, 3015–3025.

Si, Z., Guan, X., Teng, X., Peng, X., Wan, Z., Li, Q., Chen, G. and Tan, J. (2020) Identification of CYP46A1 as a new regulator of lipid metabolism through CRISPR based whole genome screening. FASEB J. 34, 13776–13791.

Solberg, L.C., Valdar, W., Gauguier, D., Nunez, G., Taylor, A., Burnett, S., Arboledas Hita, C., Hernandez Pliego, P., Davidson, S., Burns, P., Bhattacharya, S., Hough, T., Higgs, D., Klenerman, P., Cookson, W.O., Zhang, Y., Deacon, R.M., Rawlins, J.N., Mott, R. and Flint, J. (2006) A protocol for high throughput phenotyping, suitable for quantitative trait analysis in mice. Mamm. Genome 17, 129–146.

Steenbergen, R.H., Joyce, M.A., Lund, G., Lewis, J., Chen, R., Barsby, N., Douglas, D., Zhu, L.F., Tyrrell, D.L. and Kneteman, N.M. (2010) Lipoprotein profiles in SCID/uPA mice transplanted with human hepatocytes become human like and correlate with HCV infection success. Am. J. Physiol. Gastrointest. Liver Physiol. 299, G844–G854.

Stokkan, K.A., Yamazaki, S., Tei, H., Sakaki, Y. and Menaker, M. (2001) Entrainment of the circadian clock in the liver by feeding. Science 291, 490–493.

Strandberg, T.E., Kovanen, P.T., Lloyd Jones, D.M., Raal, F.J., Santos, R.D. and Watts, G.F. (2024) Drugs for dyslipidaemia: the legacy effect of the Scandinavian Simvastatin Survival Study (4S). Lancet 404, 2462–2475.

Subramanian, A., Tamayo, P., Mootha, V.K., Mukherjee, S., Ebert, B.L., Gillette, M.A., Paulovich, A., Pomeroy, S.L., Golub, T.R., Lander, E.S. and Mesirov, J.P. (2005) Gene set enrichment analysis: a knowledge based approach for interpreting genome wide expression profiles. Proc. Natl. Acad. Sci. U.S.A. 102, 15545–15550.

Sumara, G., Sumara, O., Kim, J.K. and Karsenty, G. (2012) Gut derived serotonin is a multifunctional determinant to fasting adaptation. Cell Metab. 16, 588–600.

Talreja, O., Kerndt, C.C. and Cassagnol, M. (2023) Simvastatin. In StatPearls. StatPearls Publishing.

Turek, F.W., et al. (2005) Obesity and metabolic syndrome in circadian Clock mutant mice. Science 308, 1043–1045.

Vollmers, C., Gill, S., DiTacchio, L., Pulivarthy, S.R., Le, H.D. and Panda, S. (2009) Time of feeding and the intrinsic circadian clock drive rhythms in hepatic gene expression. Proc. Natl. Acad. Sci. U.S.A. 106, 21453–21458.

Wang, Y., Liu, L. and Wang, Y. (2023) Effect of statin administration time on lipid levels and cardiovascular outcomes: a systematic review and meta analysis. Front. Pharmacol. 14, 1123456.

Whitfield, L.R., Porcari, A.R. and Alvey, C.W. (2000) Effect of dosing time on the pharmacokinetics and pharmacodynamics of atorvastatin. J. Clin. Pharmacol. 40, 758–764.

Xie, Z., Bailey, A., Kuleshov, M.V., Clarke, D.J.B., Evangelista, J.E., Jenkins, S.L., Lachmann, A., Wojciechowicz, M.L., Kropiwnicki, E. and Jagodnik, K.M., et al. (2021) Gene set knowledge discovery with Enrichr. Curr. Protoc. 1, e90.

Zayed, M.A., Jin, X., Yang, C., Belaygorod, L., Engel, C., Desai, K., Harroun, N., Saffaf, O., Patterson, B.W., Hsu, F.F. and Semenkovich, C.F. (2021) CEPT1 mediated phospholipogenesis regulates endothelial cell function and ischemia induced angiogenesis through PPARα. Diabetes 70, 549–561.

Zhang, P., Csaki, L.S., Ronquillo, E., Baufeld, L.J., Lin, J.Y., Gutierrez, A., Dwyer, J.R., Brindley, D.N., Fong, L.G., Tontonoz, P., Young, S.G. and Reue, K. (2019) Lipin 2/3 phosphatidic acid phosphatases maintain phospholipid homeostasis to regulate chylomicron synthesis. J. Clin. Invest. 129, 281–295.

Zhang, P., Munier, J.J., Wiese, C.B., Vergnes, L., Link, J.C., Abbasi, F., Ronquillo, E., Scheker, K., Muñoz, A. and Kuang, Y.L., et al. (2024) X chromosome dosage drives statin induced dysglycemia and mitochondrial dysfunction. Nat. Commun. 15, 5571.

Zimmermann, F., Roessler, J., Schmidt, D., Jasina, A., Schumann, P., Gast, M., Poller, W., Leistner, D., Giral, H. and Kränkel, N., et al. (2020) Impact of the gut microbiota on atorvastatin mediated effects on blood lipids. J. Clin. Med. 9, 1596.

